# Deciphering the postsynaptic calcium-mediated energy homeostasis through mitochondria-endoplasmic reticulum contact sites using systems modeling

**DOI:** 10.1101/2020.09.12.294827

**Authors:** A. Leung, D. Ohadi, G. Pekkurnaz, P. Rangamani

## Abstract

Spatiotemporal compartmentation of calcium dynamics is critical for neuronal function, particularly in post-synaptic spines. This exquisite level of Ca^2+^ compartmentalization is achieved through the storage and release of Ca^2+^ from various intracellular organelles particularly the endoplasmic reticulum (ER) and the mitochondria. Mitochondria and ER are established storage organelles controlling Ca^2+^ dynamics in neurons. Mitochondria also generate a majority of energy used within postsynaptic spines to support the downstream events associated with neuronal stimulus. Recently, high resolution microscopy has unveiled direct contact sites between the ER and the mitochondria, which directly channel Ca^2+^ release from the ER into the mitochondrial membrane. In this study, we develop a computational 3D reaction-diffusion model to investigate the role of MERCs in regulating Ca^2+^ and ATP dynamics. This spatiotemporal model accounts for Ca^2+^ oscillations initiated by glutamate stimulus of metabotropic and ionotropic glutamate receptors and Ca^2+^ changes in four different compartments: cytosol, ER, mitochondria, and the MERC microdomain. Our simulations predict that the organization of these organelles and differential distribution of key Ca^2+^ channels such as IP_3_ receptor and ryanodine receptor modulate Ca^2+^ dynamics in response to different stimuli. We further show that the crosstalk between geometry (mitochondria and MERC) and metabolic parameters (cytosolic ATP hydrolysis, ATP generation) influences the cellular energy state. Our findings shed light on the importance of organelle interactions in predicting Ca^2+^ dynamics in synaptic signaling. Overall, our model predicts that a combination of MERC linkage and mitochondria size is necessary for optimal ATP production in the cytosol.

## 1 Introduction

Compartmentalized calcium handling in postsynaptic structures underlies synaptic communication and controls synaptic plasticity, which is the bidirectional modulation of synaptic strength that is thought to underlie learning and memory formation [1]. In excitatory neurons during synaptic transmission, glutamate released from the presynaptic bouton leads to a localized increase in calcium at the postsynaptic dendrite that is critical for the induction of synaptic plasticity [2, 3]. At the postsynaptic site, small bulbous protrusions called dendritic spines act as biochemical computation units that regulate the duration and spread of postsynaptic calcium fluxes produced by glutamatergic neurotransmission [4]. The temporal dynamics of calcium, and the coupling to downstream signaling pathways in dendritic spines depend on many factors, including the nature of stimulus as well as the positioning of the calcium storage organelles, the endoplasmic reticulum (ER) and the mitochondria [5, 6]. A continuous tubular network of ER spreads throughout the dendrites and extends into the spines either as a simple smooth tubule or a spine apparatus [7–9]. Mitochondria of various lengths occupy a major portion of dendritic branches and associate closely with the ER particularly at the dendritic base of spines [10–12]. The intimate contact between mitochondria and ER along dendrites allows for a functional inter-organellar coupling and plays a central role in the regulation of the postsynaptic calcium dynamics (Figure 1a). Dysregulation of mitochondria and ER coupling have been demonstrated in neurodegenerative diseases such as Alzheimer’s [13–15] and Parkinson’s [15, 16], although the underlying mechanisms are yet to be elucidated.

**Figure 1:**
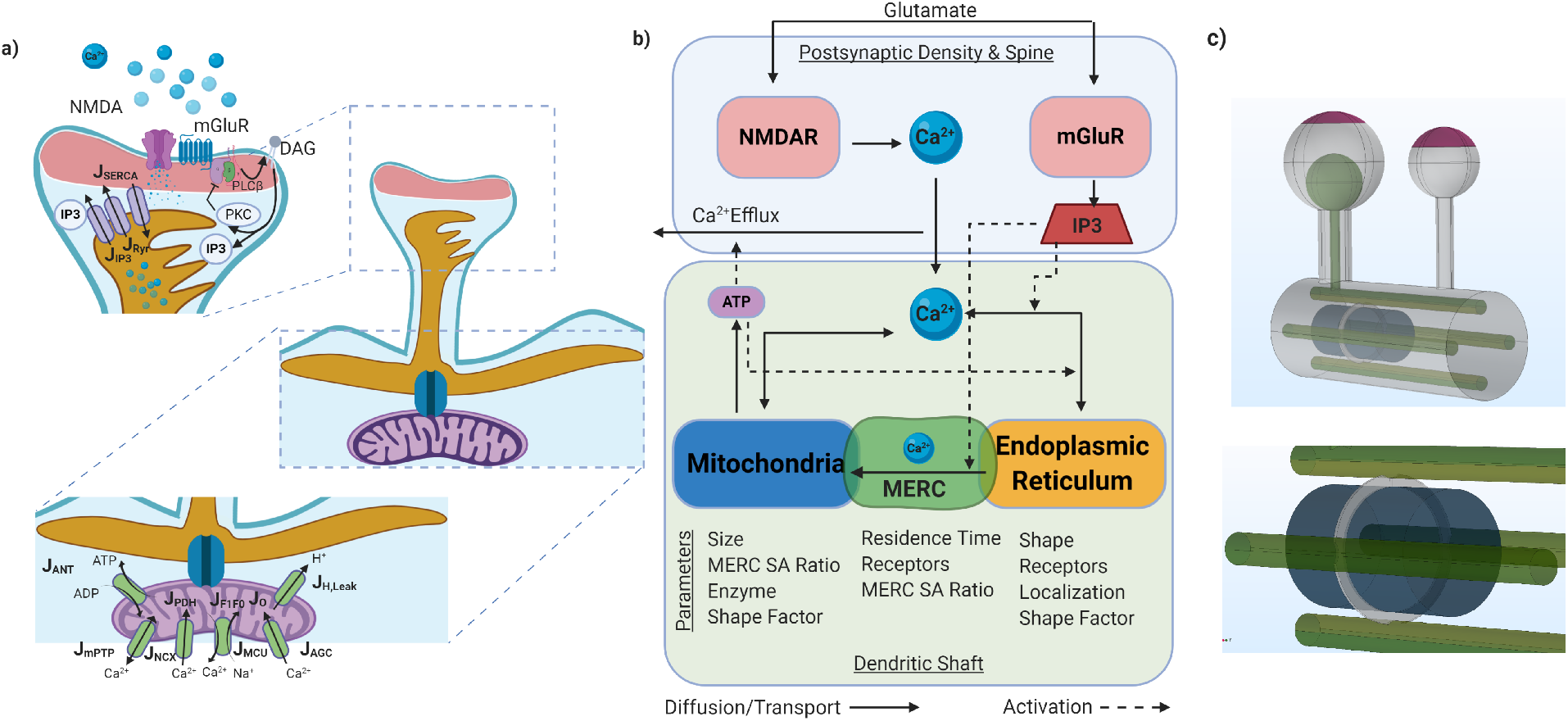
Schematic of the model architecture and fluxes in each compartment. a) Glutamatergic signaling in a postsynaptic spine leads to an influx of Ca^2+^ from the extracellular space and from the endoplasmic reticulum. Dynamics of Ca^2+^ have a direct impact on mitochondrial function that alters the ATP production. [25, 28]. The main fluxes involved in the model are shown here and the mathematical descriptions are given in Table S2. b) The main compartments and investigated parameters in the model are shown in this schematic. In this figure, solid lines represent diffusive or transport fluxes. Dashed lines represent an activation of fluxes. c) Model geometry for a dendritic spine developed using COMSOL Multiphysics^®^. Two spines, one with and one with the spine apparatus are connected by a dendritic shaft containing the ER and mitochondria. PSD, Purple; ER, Green; MERC, Grey; Mitochondria, Blue. SA Ratio refers to the ratio of surface area of MERCs to total mitochondria surface area

Physical contact sites between mitochondria and ER (Mitochondria-ER Contact Sites, or MERCs) were first observed by electron microscopy, then confirmed by dimerization-dependent fluorescent proteins [17, 18]. MERCs provide a direct avenue for calcium to transfer from the ER to the mitochondria [19]. Because the concentration of ER calcium is several orders of magnitude higher than cytosolic calcium, the existence of these contact sites can result in a direct conduit for increasing mitochondria calcium concentration. These MERCs have been observed to be 2-20% of a mitochondria’s surface area [17]. MERCs are also essential for mitochondrial functions and ATP production [20]. While the exact mechanism for the formation of these sites is unclear, there is evidence that binding proteins link via calcium channels on the surface of the respective organelle [18, 21]. The role of organelles and MERC in calcium dynamics of dendritic spines is not yet fully understood.

Computational modeling has played a pivotal role in providing insight into calcium dynamics in neurons [22–24]. Bertram et al. [25], Han et al. [26], Wacquier et al. [27] and others have created models addressing the role of calcium in mitochondria. Bertram et al. [25] simplified a mathematical model for mitochondrial calcium dynamics in pancreatic beta cells originally developed by Magnus et al. [28] to explain experimentally derived results [29] in which calcium increases NADH under low glucose and reduces NADH in high glucose. Han et al. [26] explicitly modeled the role MERC in a pancreatic beta cell in a compartmental model for the case of a healthy and diabetic cells and predicted a connection between obesity, MERCs, and calcium dynamics. Qi et al. similarly modeled mitochondrial-ER calcium flux into the mitochondria as a function of distance between IP_3_R and MCU to predict an optimal ER-to-mitochondria distance for the regulation of mitochondrial calcium [30]. However, none of these models are specific to unique circumstances of a dendritic spine or neuronal stimulus patterns. Previously, we and others have shown that spatial modeling of signaling pathways can provide insight into how cell shape and organelle organization can affect the spatiotemporal dynamics of second messengers and signaling cascades [31–36] Although spatial models for neuronal calcium signaling models exist [**?**, 37, 38], these do not focus on the metabolic consequences of calcium and incorporate mitochondria.

In this study, we investigated the role of the ER, mitochondria, and MERC in modulating the spatiotemporal dynamics of calcium and localized ATP production in postsynaptic spines using computational modeling. We specifically sought to answer the following questions: What are the spatiotemporal dynamics of Ca^2+^ and IP_3_, in response to a glutamate stimulus, in the spine head, spine shaft, and the mitochondria? How does the presence or absence of MERCs affect the dynamics of these second messengers and alter the energy landscape in these locations? And finally, how do different geometric features such as mitochondrial length and MERC surface area ratio affect Ca^2+^ handling and energy production in spines? To answer these questions, we developed a spatiotemporal model of Ca^2+^ dynamics in a portion of a dendrite with spatially resolved ER and mitochondria in idealized geometries. We found that the MERC leads to a significant increase in mitochondrial calcium, and subsequently ATP production. Additionally, increasing the surface area of the MERC, increases the calcium influx into the mitochondria, providing insight into how the different extent of MERC can give rise to different calcium states. Finally, we predict metabolic parameters, such as cytosolic hydrolysis, nucleotide transport from the mitochondria to the cytoplasm, and rate of ATP synthesis to be key deciding factors in the delicate balance of calcium signaling and energy homeostasis in the postsynaptic spine.

## Results

We constructed a spatial model with five compartments: the postsynaptic density (PSD), the cytosol, one mitochondria, the ER, and a mitochondria ER Contact region (MERC) (Figure 1a, b). In our simplified geometry (shown in Figure 1c), the dendritic spine is modeled as a sphere attached to the large cylindrical dendritic shaft by a narrow cylindrical spine neck, geometrically defined in Table S1. Although the morphology of a spine has been shown to govern the magnitude and stability of calcium transients in previous studies [24, 31], in this study we simplify complex spine morphology to idealized geometries to focus on the role of mitochondria in neuronal calcium dynamics. While the interconnected ER tublues are widely distributed throughout the cytoplasm, we approximate the ER within the dendritic shaft as long cylinders, keeping ER to cytoplasmic volume ratios constant. The mitochondrion was modeled as a wide cylinder in the center of these four ER. Although dendritic mitochondria have a wide range of sizes from sub micron to to more than 10 μm with complex inner membrane and outer membrane morphologies [39], we approximate a relatively small mitochondria of 0.6 μm localized at the base of a dendritic spine. The sizes of the compartments in the model are given in Table S1. The details of the model development and numerical methods are given in the supplementary material and in Table S3

### Spatial dynamics of Ca^2+^ and IP_3_ in spines with and without MERC

We conducted simulations with a 1 Hz stimulus of glutamate for 5 s and ran the simulations until steady state. We also investigated how the two different glutamate receptors, NMDAR and mGluR contributed to the effect of signaling dynamics in the presence and absence of MERC (Figure 2). We dissected these effects by comparing the contribution of NMDAR only, mGluR only, and when both receptors are active. In all simulations, only the larger spine with the spine apparatus was stimulated with the 1 Hz pulse train. In Figure S1, we instead stimulate the neighboring smaller spine and find minimal differences in dynamics. We conducted these simulations in two geometries – one without the MERCs (no gray ring) and one with MERC consisting of 10% of the total surface area (Figure 1c). Spatial maps at 1 s and 5.1 s were chosen to demonstrate the diffusivity of the Ca^2+^ and IP_3_ dynamics (shown in Figure 2). Corresponding temporal dynamics at different locations are shown in Figure S2.

**Figure 2:**
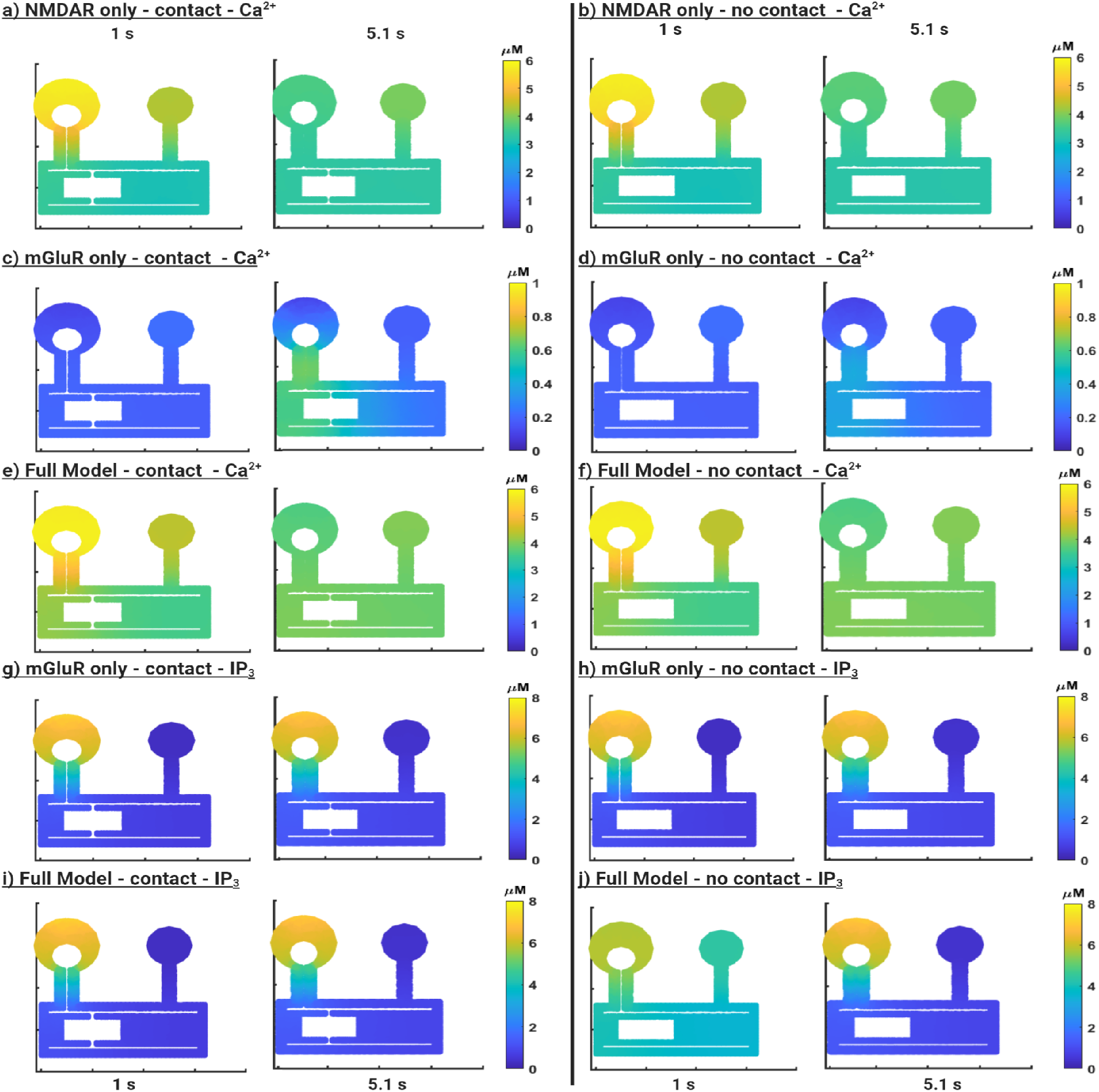
Receptor-based simulations of glutamatergic signaling in spatial calcium model. Finite element simulations of dendritic spines including mitochondria, ER, and mitochondrial-ER contacts(MERC). **a**) Ca^2+^ dynamics in the stimulated and neighboring spine when only NMDAR are stimulated in the presence of MERC. **b**) Same as (**a**) in the absence of MERC. In this case, there is no production of IP_3_. **c**) Ca^2+^ dynamics when only mGluR is stimulated in the presence of MERC. **d**) Same as (**c**) in the absence of MERC. **e**) Ca^2+^ dynamics in the stimulated and neighboring spine when both NMDAR and mGluR are stimulated in the presence of MERC. **f**) Same as (**e**) in the absence of MERC. **g**) IP_3_ dynamics in the stimulated and neighboring spines when only mGluR is stimulated in the presence of MERC. **h**) Same as (**g**) in the absence of MERC. **i**) IP_3_ dynamics in the stimulated and neighboring spines when both NMDAR and mGluR is stimulated in the presence of MERC. **j**) Same as (**i**) in the absence of MERC.

We first looked at Ca^2+^ dynamics and IP_3_ distribution with and without MERC (Figure 2). In the presence of MERC, we note that when only NMDAR is active, the calcium dynamics are quite rapid in the stimulated spine and the neighboring spine (Figure 2a). The peak calcium concentration is higher in the spine head (6 μM) and about half that in the dendritic shaft (3 μM) proximal to the stimulated spine. In contrast, the neighboring spine has a peak Ca^2+^ concentration of about 4 μM with a distinctly lower gradient between spine head and shaft calcium. These results follow from the fact that synaptic activation of NMDAR located at the PSD leads to an increased Ca^2+^ influx and therefore, the stimulated spine has the peak calcium concentration. When we look at the simulations without MERC (Figure 2b), we note that the absence of MERC has negligible effects on the spine Ca^2+^ concentration dynamics in both the stimulated and neighboring spines. This is predicted NMDAR affects rapid Ca^2+^ influx from the spine head and is not immediately affected by organelle calcium dynamics.

When we consider the effect of mGluR response only, we first note that the calcium response is delayed when compared to the stimulus and the NMDAR response in both the stimulated spine and the neighboring spine (Figure 2c). This is because Ca^2+^ release in this scenario is primarily IP_3_-mediated and Ca^2+^ is released from the spine apparatus and the ER. Since the mGluR-only system does not have an influx of calcium, the average concentration of calcium is significantly lower (1 μM). Comparing the scenario with and without MERC, we note that the MERC compartment essentially allows for an increased and rapid early release of Ca^2+^ from the base of the stimulated spine (Figure 2c,d) than in the case without MERC. When we look at the corresponding IP_3_ dynamics in the stimulated spine (Figure 2g,h), we note that the immediate response of the stimulated spine, the spatial gradient of IP_3_, primarily from the spine head to the dendrite, consistent with the mGluR location and PIP_2_ hydrolysis from the plasma membrane. The neighboring spine receives IP_3_ only through diffusion and therefore has a lower concentration and no observable gradients of IP_3_. When there is no MERC, IP_3_ concentrations are similar to the case with MERC because IP_3_ is an early response to mGluR and not affected at these times by Ca^2+^. In both cases, the neighboring spine shows delayed IP_3_ dynamics and reduced concentrations.

When both stimuli are included, it becomes immediately obvious that NMDAR contributes dominantly to Ca^2+^ dynamics (Figure 2e,f) and while IP_3_ dynamics are dominated by the mGluR activation (Figure 2i,j). We also note that there are spatial differences in the Ca^2+^ dynamics in the stimulated spine when both NMDAR and mGluR are present as opposed to the individual stimuli because now the Ca^2+^ influx is both from the PSD and from the spine apparatus.

We also applied no stimulus and found the system quickly approaches an equilibrium for Ca^2+^, although ATP decay on a longer timescale (Figure S3), discussed in detail below. To further characterize the cytosolic Ca^2+^ dynamics in response to stimuli in this system, we changed the frequency of the stimulus comparing 1, 10, and 50 Hz stimulus over 5 seconds (Figure S4). We found that Ca^2+^ in the stimulated and neighboring spines showed a frequency dependence for the 1 and 10 Hz input and an integrated calcium behavior (reading only the lower frequencies for the 50 Hz) (Figure S4a, b).

### Presence of MERCs alter organelle Ca^2+^ dynamics and ATP production

Next, we investigated the effect of MERCs on Ca^2+^ dynamics in the ER and mitochondria with the same simulation set as in Figure 2. In the dendrite, the ER acts as an intracellular calcium storage compartment and surface receptors on the ER, such as ryanodine and inositol 1,4,5-trisphosphate (IP_3_R), rapidly release calcium into the cytosol [40]. Sacroplasmic/endoplasmic reticulum Ca^2+^-ATPases (SERCAs) can result in high uptake of Ca^2+^ into the ER (> 300 μM [41]) through the consumption of ATP. Ca^2+^ is then released from the ER through IP_3_R and ryanodine receptors (RyRs) [42]. Ca^2+^ regulation and mitochondrial ATP production are crucial for synaptic function and neuroplasticity [43]; mitochondria in synaptic terminals aid neurotransmission by producing ATP, buffering calcium, and local protein translation [10, 44].

In our simulations, when MERCs are present, we observed that the mitochondrial calcium is significantly higher for all three receptor conditions when compared to the case without MERCs (Figure 3a). This effect was larger when only NMDAR are activated relative to the mGluR or full model. Because MERCs increase the total influx of calcium into the mitochondria, while the efflux is limited by cytosolic exchange, there is a notable accumulation of calcium within the mitochondria. Comparison of mitochondrial calcium dynamics with and without MERCs reveals that the presence of MERCs smooths out the frequency of the glutamate stimulus and time of peak Ca^2+^ concentration is slightly delayed when compared to the scenario without MERCs. These changes in the calcium dynamics can be better understood by comparing the integrated Ca^2+^ (area under the curve, AUC), which reveals that the mitochondrial Ca^2+^ AUC is higher in the presence of MERCs. This is the first indication from our simulations that spatial compartments such as MERCs can alter Ca^2+^ dynamics in organelles, consistent with other hypotheses in the literature [17, 45, 46]. On the other hand, no dramatic differences were observed in the ER Ca^2+^ dynamics with and without MERC (Figure 3b). Although the MERC is directly connected to the ER and there is significant calcium flux from the ER to the MERC, the relative size of the ER keeps the overall concentration of calcium constant in both cases.

**Figure 3:**
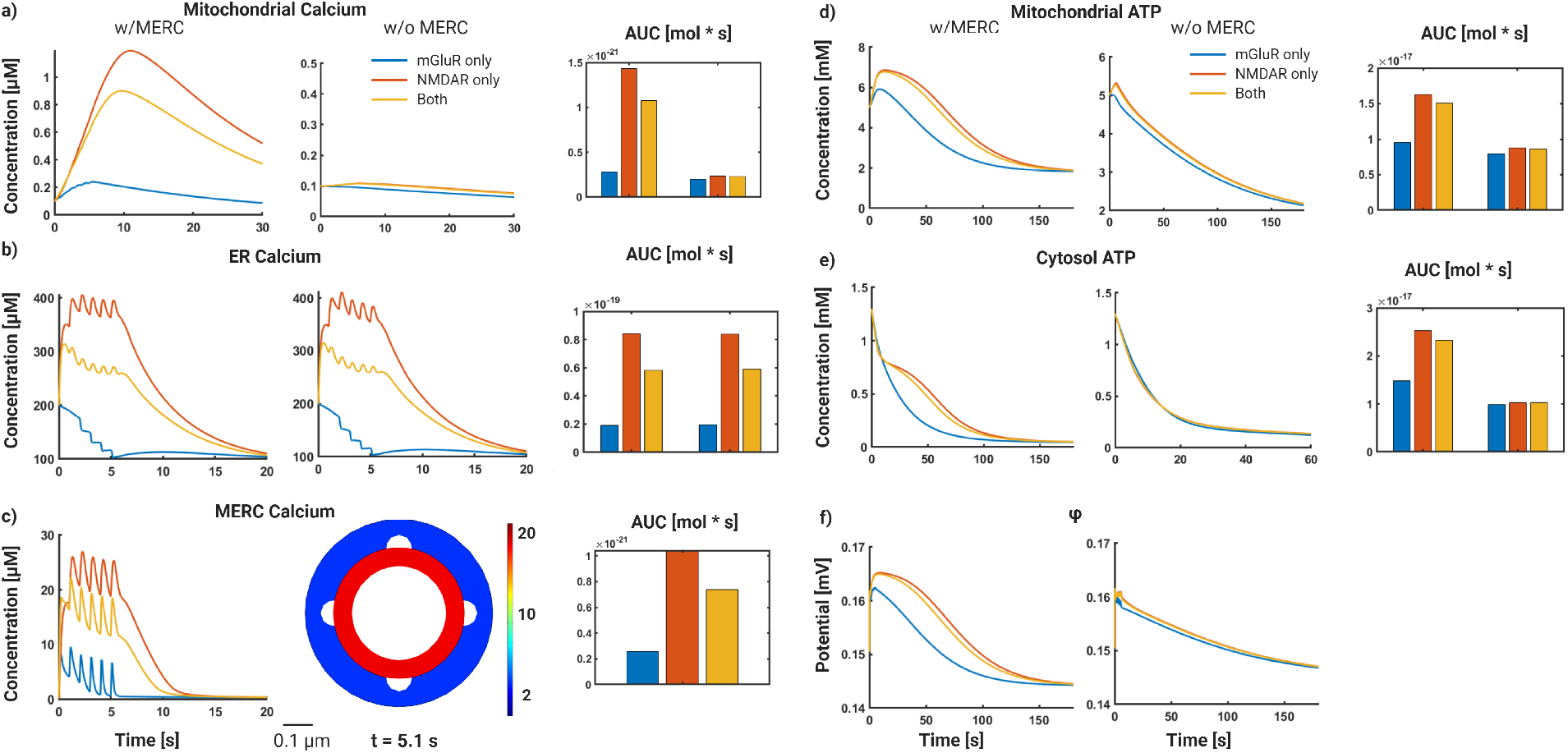
Receptor-based simulations of glutamatergic signaling in spatial calcium model. Finite element simulations of dendritic spines including mitochondria, ER, and mitochondrial-ER contacts(MERC). In all the plots, the blue line is for mGluR only, red for NMDAR only and yellow for both receptors. **a**) Mitochondrial Ca^2+^ dynamics in the presence (left) and absence (middle) of MERC. Total Ca^2+^ calculated using the area under the curve (AUC) (right) **b**) ER Calcium dynamics with (left) and without MERC (middle); AUC (right). **c**) Dynamics of MERC Ca^2+^ in the MERC case only (left); cross section of MERC ring showing a higher concentration of Ca^2+^ in the MERC ring as opposed to the general cytoplasm (right). **d**) Dynamics of mitochondrial ATP; with MERC (left), without MERC (middle), AUC (right). **e**) Dynamics of cytosolic ATP; with MERC (left), without MERC (middle), AUC (right). **f**) Mitochondrial membrane potential; with MERC (left), without MERC (right).

Perhaps the most interesting feature of this model is that MERC act as a Ca^2+^ microdomain by localizing high Ca^2+^ concentrations for a finite duration in a receptor dependent manner (Figure 3c). Thus, our model predicts that the presence of MERC plays an important role in mitochondrial and MERC Ca^2+^ dynamics. As a result, we see an impact on mitochondrial and cytosolic ATP dynamics (Figure 3d,e). In the presence of MERC, both mitochondrial and cytosolic ATP are increased and and have prolonged dynamics. The increased production and prolonged dynamics is dependent on the calcium diffusion within the system. In Figure S5, we show that decreasing the diffusion coefficient leads to a delay in mitochondrial calcium peak, with a corresponding increase in cytosolic and mitochondrial ATP. We also note that compared to the Ca^2+^ dynamics in the ER and the MERC, the mitochondrial Ca^2+^ and ATP dynamics are smoother, indicating that the mitochondria retain the lower frequency information and not the higher frequency. When we compare the mitochondrial ATP, we note that the presence of MERCs can substantially increase mitochondrial ATP production (Figure 3d) and cytosolic ATP availability (Figure 3e). The dynamics of the mitochondrial membrane potential (Ψ_*m*_) are also altered in the presence of MERCs (Figure 3f), consistent with the mitochondrial ATP dynamics. Interestingly, when we investigated the role of different stimulus frequencies, we found that organelle Ca^2+^ and ATP dynamics were more sensitive to the change from 1 to 10 Hz than from the 10 to 50 Hz (Figure S4c-f). We note that varying the diffusion coefficient of Ca^2+^ did not alter the results from our model significantly (Figure S5).

Thus, the MERC compartment, which is technically part of the cytosol, is enriched in calcium even though there is no physical membrane separating the two compartments. By serving as a direct conduit for Ca^2+^ influx from the cytosol, we suggest that MERCs play an important role in localizing Ca^2+^ and thereby increasing ATP production locally at active synapses.

### Effect of mitochondrial size and MERC area fraction on Ca^2+^ and ATP dynamics

The mitochondria present in dendrites of neurons are abundant and can vary in size [10, 39]. Separately, the area fraction of MERCs are also known to vary [17]. Do these physical variables have an impact on the production of ATP in the mitochondria? To address this question, we conducted two sets of simulations: vary mitochondrial length while maintaining MERC area fraction, and (ii) vary MERC area fraction in mitochondria of 3 sizes (Figures 4 and 5, respectively).

**Figure 4:**
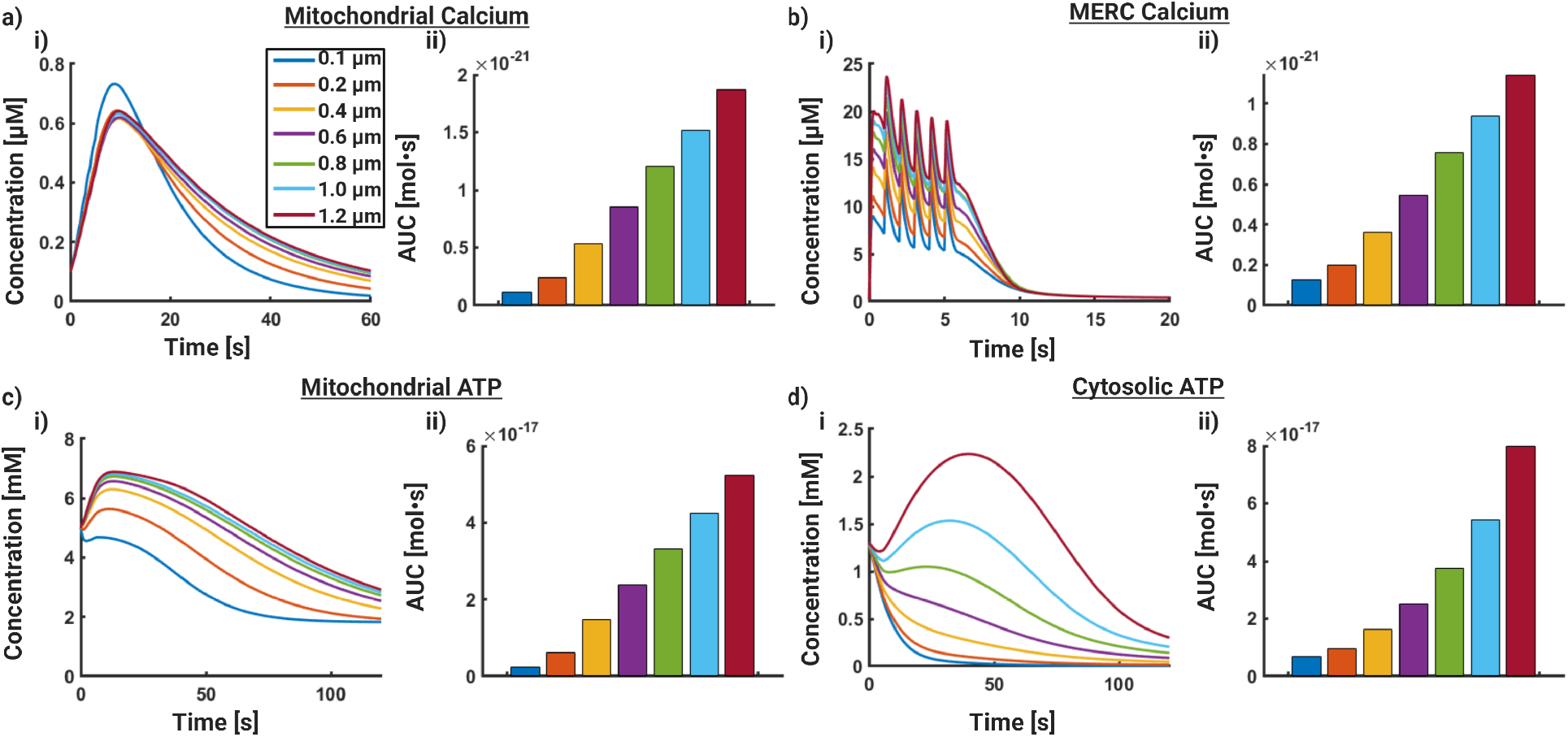
Mitochondrial length increases the magnitude of mitochondrial Ca^2+^ and ATP. In these simulations we increased the length of the mitochondria while keeping the MERC area fraction constant at 0.1. **a**) (i) Volumetric average dynamics of mitochondrial Ca^2+^ increases with increasing mitochondrial length; AUC for mitochondrial Ca^2+^, integrated over time and mitochondrial volume. **b**) (i) Volumetric average dynamics of MERC Ca^2+^; (ii) AUC for MERC Ca^2+^. **c**) (i) Volumetric average dynamics of mitochondrial ATP increases with increasing mitochondrial length; (ii) AUC for mitochondrial ATP, integrated over time and mitochondrial volume. **d**) (i) Volumetric average dynamics of cytosolic Ca^2+^ increases with increasing mitochondrial length; (ii) AUC for mitochondrial Ca^2+^, integrated over time and mitochondrial volume.

**Figure 5:**
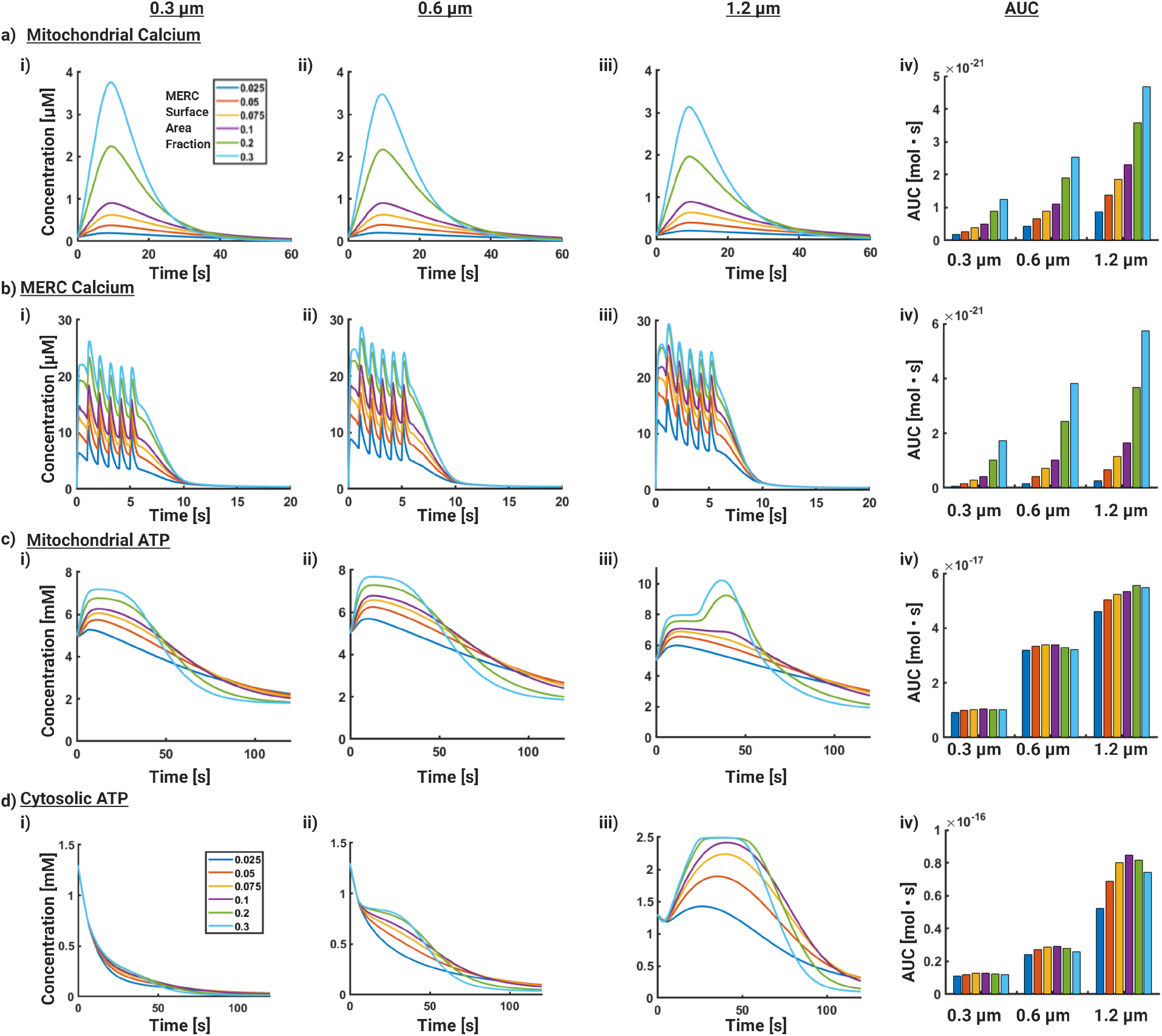
Effect of MERC area fraction on ATP production. Here we vary the size of the mitochondria and the area fraction of MERC systematically. Three different mitochondrial sizes (0.3 *μm*, 0.6 *μm*, and 1.2 *μm*) and six different MERC area fractions ranging from 0.025 to 0.3 were simulated. **a**) Mitochondrial Ca^2+^ dynamics and AUC for the different cases. **b**) MERC Ca^2+^ dynamics and AUC for the different cases. **c**) Mitochondrial ATP dynamics and AUC for the different cases. **d**) Cytosolic ATP dynamics and AUC for the different cases.

We first explore the effect of mitochondrial size on organelle Ca^2+^ and ATP dynamics (Figure 4). In these simulations, we maintained the MERC area fraction (calculated as the ratio of MERC surface area in contact with mitochondria to the total mitochondrial surface area) as 0.1 while varying the mitochondrial size from 0.2 to 1.2 *μm*. Since the geometries have different volumes, we calculate the AUC as an integral over both time and volume to show the total number of moles of the species integrated over time.

Our simulations showed that increasing mitochondrial size clearly favored increases in mitochondrial Ca^2+^ (Figure 4a) and MERC Ca^2+^ (Figure 4b). We note that the increase in the Ca^2+^ begins to saturate (see peak Ca^2+^ values in Figure 4a) after a certain size of mitochondria suggesting that there is no additional advantage of large mitochondria in terms of Ca^2+^ concentration. This effect is also seen in the MERC Ca^2+^. Correspondingly, we observed increases in mitochondrial ATP production capacity but also a saturation at higher mitochondrial sizes (Figure 4c). Thus far, our results are consistent with our expectation that increasing mitochondrial size increases the total amount of Ca^2+^ and ATP produced in the mitochondria. In this model, mitochondrial size is limited by our dendrite segment size, however our results suggest that a larger mitochondria would produce more ATP.

When we look at the cytosolic ATP, we note that smaller mitochondria actually show a decrease in cytosolic ATP. As the mitochondrial size increases, cytosolic ATP increases and at larger mitochondrial sizes the kinetics are prolonged (Figure 4d). This effect of mitochondrial size on cytosolic ATP can be understood as a result of the competition between the boundary fluxes on the mitochondrial membrane for ANT and the hydrolysis rate occurring in the cytosol. As the mitochondrial size increases, the corresponding surface area increases, increasing the flux across the mitochondrial membrane into the cytosol. This change in flux also alters the kinetics of cytosolic ATP. Increasing the length of the mitochondria increases the duration of Ca^2+^ flux into the mitochondria, especially when there is sufficient MERC to facilitate Ca^2+^ influx. This increase in the flux shifts the overall Ca^2+^ concentration, resulting in a longer half-life of mitochondrial ATP and an increase in peak times for cytosolic ATP (Table S4).

Thus, our model predicts that the presence of MERCs alone does not inherently confer an advantage for increased cytosolic ATP for all mitochondrial sizes. Rather, an equilibrium between rapid ATP production in the mitochondria through MERCs and a rapid ATP delivery to the cytosol through ANT are required to balance ATP production and supply in dendrites. ANT also drives ADP transport between the cytosol and mitochondria, which is a necessary substrate for ATP production. For effective ATP availability in the dendrites our model predicts two requirements – production of ATP in the mitochondria and availability of ATP in the cytosol. Production of ATP in the mitochondria depends on MERC area fraction and mitochondrial size. Availability of ATP in the cytosol depends on the flux of ATP from the mitochondria through the ANT and the consumption of ATP through a lumped hydrolysis rate.

### Combination of increased mitochondrial size and MERC area fraction buffers ATP production capacity

We next investigated the role of MERC and mitochondrial sizes in governing ATP production capacity. Since MERCs provide a direct conduit to increase mitochondrial Ca^2+^, we varied the area fraction of MERCs with mitochondria of different sizes (0.3, 0.6, and 1.2 *μ*m). These lengths are supported by observations that Ca-CaMKII activity can decrease mitochondrial length over a range of synaptic activity levels [47]. Particularly, we sought to find if increasing MERC area fraction could confer an advantage to smaller mitochondria, compensating for volumetric disadvantage with increased calcium. Experimental evidence indicates that MERCs range from 2-20 % of mitochondria surface area depending on the metabolic state and type of cell [45, 48, 49]. We conducted simulations in small (0.3 μm), medium (0.6 μm), and large mitochondria (1.2 μm), with a range of MERC area fractions ranging from 2.5 % to 30 % (Figure 5) for a fixed value of the ATP consumption rate of 200 μM/s. We note that, while area fraction ranges have been experimentally quantified, they need not necessarily form a ring-like structure that we model here [17, 48, 50, 51]. As shown in Figure S6, the effect of MERCs on cytosolic Ca^2+^ and IP_3_ are not dramatically affected by overall size of MERCs.

We observe that increasing the MERC area fraction increases mitochondrial Ca^2+^ across all sizes but the peak concentration of Ca^2+^ decreases with increasing mitochondrial size (Figure 5a). This effect is true for both the small and large mitochondria, and the larger mitochondria have higher calcium concentration and total amounts (by virtue of larger volumes). MERC Ca^2+^ dynamics also show an increase with increasing MERC area fraction for all mitochondrial sizes with a proportionally small increase in the peak Ca^2+^ (Figure 5b). Mitochondrial ATP production increases with the MERC area fraction within a given mitochondrial size and also increases with larger mitochondrial sizes (Figure 5c). Interestingly, the dynamics of mitochondrial ATP show a faster decay with increasing MERC area fraction for small and medium mitochondria (Figure 5c, i,ii). For the large mitochondria, we observe a second and prolonged peak of mitochondrial ATP as the area fraction of MERC increases (Figure 5c,iii). Thus the combination of MERC area fraction and mitochondrial sizes results in synergistic effects on mitochondrial ATP production. However, looking at the cytosolic ATP dynamics, we note that not all mitochondrial sizes or MERC fractions are conducive to increased ATP availability (Figure 5d). Small mitochondria show practically no increase in cytosolic ATP and are unaffected by MERC area fraction (Figure 5d, i). Medium size mitochondria begin to show some response in that increasing the MERC area fraction prolongs the decay time but no dramatic increase in cytosolic ATP is observed (Figure 5d,ii). Large mitochondria show a strong sensitivity to MERC area fraction – first, large mitochondria show an increase in cytosolic ATP even for small MERC area fractions and second, increasing MERC area fraction in large mitochondria results in a larger ATP concentration in the cytosol (Figure 5iii).

### Energy availability at the synapse is a combination of mitochondrial geometry, MERC area fraction, and kinetics of ATP production and consumption

Finally, we asked the following question: how is the balance between the kinetics of ATP production through F1FO, ATP transport through ANT, and ATP consumption (modeled using a lumped hydrolysis term, K_*HY D*_) influenced by mitochondrial size and MERC area fraction? We conducted a series of simulations to map the corresponding phase spaces that hold the answer to this question. For all simulations, we varied mitochondrial size from 0.3 - 1.2 μm and MERC area fraction (defined as the ratio of MERC SA to mitochondria SA) from 0 (no contact) to 0.2. Then, we calculated the AUC for mitochondrial ATP and the AUC for cytosolic ATP as our scalar readout.

To investigate the role of the rate of ATP production by the F1,FO ATP synthase, we varied the kinetic parameter *V*_*F*1,*FO*_ (Figure 6a) (Tables S2 and S5). This variation effectively captures the changes to either the activity of the ATP synthase or the number of ATP synthases. We varied the value of *V*_*F*1*FO*_ by two orders of magnitude with respect to our control value of 35 μM. We found that as we increased *V*_*F*1*FO*_, mitochondrial ATP increases with mitochondrial size and MERC area fraction across all the values of *V*_*F*1*FO*_. For higher values of *V*_*F*1*FO*_, however, we lose the dependence on MERC area fraction obtaining simply a linear dependence on mitochondrial size. In contrast, cytosolic ATP shows a non-monotonic dependence on *V*_*F*1*FO*_ for different mitochondrial lengths and MERC SA ratio. For low *V*_*F*1*FO*_, cytosolic ATP (Figure 6b) is maximum for large mitochondrial length and for MERC area fractions close to 5%. As the MERC area fraction increases, the cytosolic ATP decreases. However, for medium values of *V*_*F*1*FO*_, cytosolic ATP increases with both mitochondrial length and MERC SA ratio, indicating the increasing rate of ATP synthesis gives high cytosolic ATP for a wide range of MERCs. Further increase in *V*_*F*1*FO*_ did not confer any additional advantage on cytosolic ATP; instead the cytosolic ATP was lower for larger mitochondrial lengths and MERC area fraction. Although the AUC captures the total production of ATP, to study the dynamics we plot a selection of points in the phase diagram in Figure S7. We show that the dynamics of ATP production in the mitochondria and ATP availability in the cytosol are dramamtically altered by the values of *V*_*F*1*FO*_ and the geometric parameters (Figure S7) suggesting a more complex spatiotemporal regulation.

**Figure 6:**
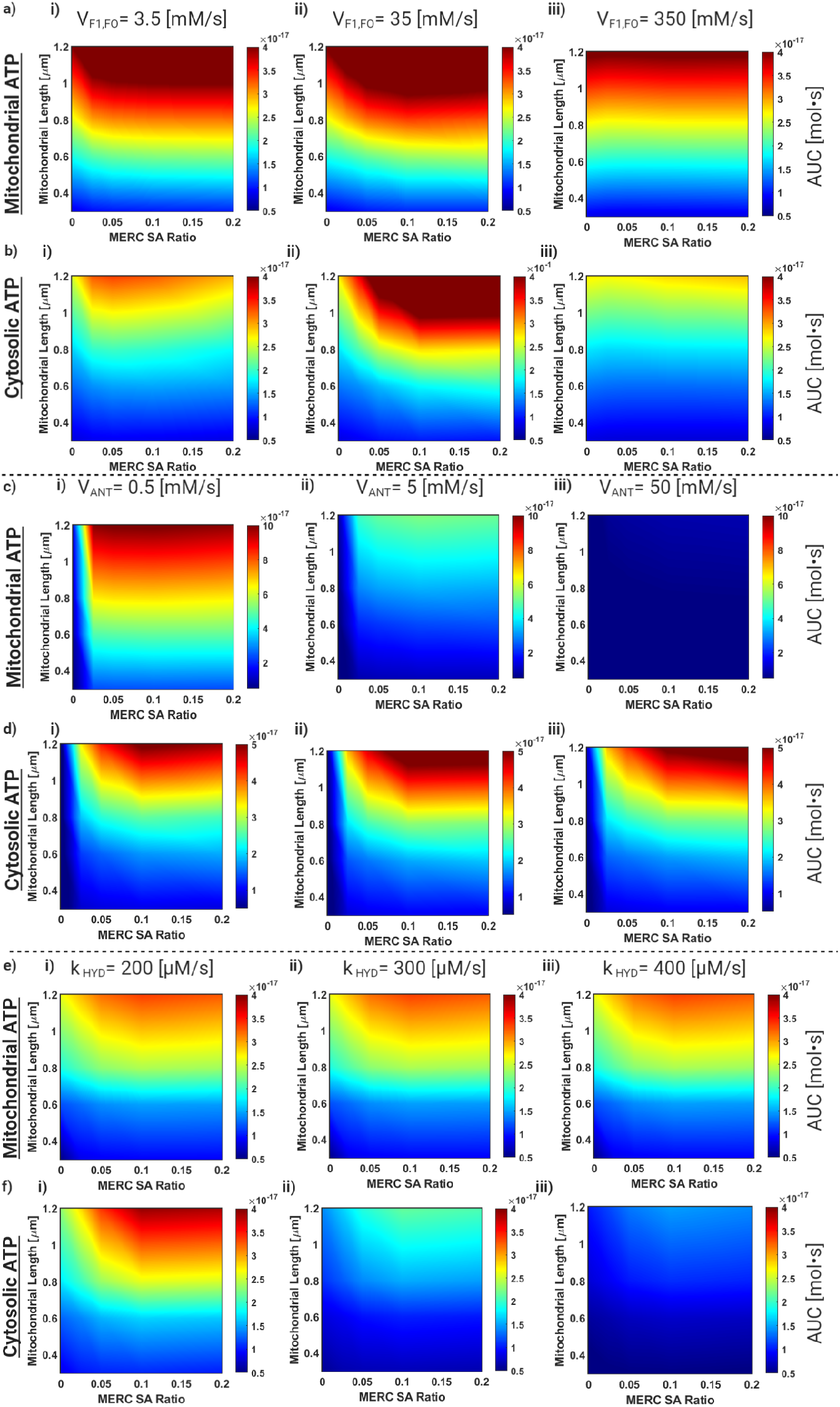
Model overview and discussion of geometric control of metabolic balance. **a**) Cytosolic ATP AUC phase map for **i**) *V*_*F*1,*FO*_ = 3.5*mM/s*, **ii)** *V*_*F*1,*FO*_ = 35*mM/s*, and **iii)** *V*_*F*1,*FO*_ = 350*mM/s*. **b**) Same as **a**) for mitochondria ATP. **c**) Cytosolic ATP AUC phase map for **i**) *V*_*ANT*_ = 0.5*mM/s*, **ii**) *V*_*ANT*_ = 5*mM/s*, **iii**) *V*_*ANT*_ = 500*mM/s*. **d**) Same as **c**) for mitochondria ATP. **e**) Cytosolic ATP AUC phase map for **i)** *K*_*HY D*_ = 200*mM/s*,**ii)** *K*_*HY D*_ = 300*mM/s*, and **iii)** *K*_*HY D*_ = 400*mM/s*. **f**) Same as **e)** for mitochondria ATP.

These results suggest that increasing *V*_*F*1*FO*_ increases mitochondrial ATP, but the impact on cytosolic ATP is mixed. This could be because either the model is incomplete because we don’t consider the morphology of the inner mitochondrial membrane (see [52] for a detailed inner mitochondrial membrane model) or that the production of ATP is not matched by the transport to the cytosol.

Next, we changed the value of *V*_*ANT*_ to study the role of ATP transport across the mitochondrial membrane on the energetic balance between mitochondria and cytosol (Figure 6c). For the low value of *V*_*ANT*_, the mitochondrial ATP AUC increases with mitochondrial length and MERC area fraction. The cytosolic ATP AUC also increases for mitochondrial length and MERC area fraction. When *V*_*ANT*_ increases, the accessible states for mitochondrial ATP AUC decrease with respect to our geometric parameters (Figure 6c,ii) but remain unchanged for cytosolic ATP(Figure 6d,ii). For high values of V_*ANT*_ we see a similar trend as the difference in mitochondria ATP AUC states is nearly abolished (Figure 6c,iii), but has little effect on the cytosolic ATP. Thus, *V*_*ANT*_ is a critical parameter in maintaining mitochondrial ATP generation rates across different mitochondrial lengths and MERC area fractions. However, V_*ANT*_ does not have an influence on the total production of ATP in the system.

Finally, in Figure 6e, we explore the role of the rate of ATP consumption in our system, by varying *k*_*hyd*_ Table S5 in the cytosol. This lumped parameter effectively captures the energy consumption in the post-synaptic spine; over 60 % of neuronal energy consumption is consumed during active calcium signaling, which occurs at the PM and ER boundaries [53]. We note that there is limited effect of changing the AUC of mitochondrial ATP across different values of *k*_*hyd*_ and MERC SA ratio. There is a linear increase in Mitochondrial ATP AUC with respect to mitochondrial length (Figure 6e). Cytosolic ATP (Figure 6f), on the other hand, decreases with increase in *k*_*hyd*_, as expected. For low values of *k*_*hyd*_, cytosolic ATP increases with mitochondrial lengths and MERC surface area. As *k*_*hyd*_ increases, the consumption of ATP essentially overrides any geometric dependence. Larger mitochondria are able to offset this higher consumption, but as the energy consumption increases the system is unable to reach higher energy states at the current geometry. Thus, our model shows that the metabolic parameters associated with ATP production, transport, and consumption in conjunction with MERC surface area ratio and mitochondrial length are important for ATP availability.

## Discussion

In this study, we have investigated how the presence of MERCs can alter Ca^2+^ and ATP dynamics in the postsynaptic spine and surrounding areas using computational modeling. Our model predicted that the presence of MERCs can lead into increased mitochondrial calcium transients and increased mitochondrial ATP production consistent with experimental observations [47, 50, 54]. We also showed that mitochondrial size and MERC area fraction play a critical role in the balance of mitochondrial ATP production and cytosolic ATP by regulating the fluxes into and outside of the mitochondria [47]. And finally, we predict that metabolic parameters associated with mitochondrial function are also important determinants of this balance between energy production and energy availability (Figure 7).

**Figure 7:**
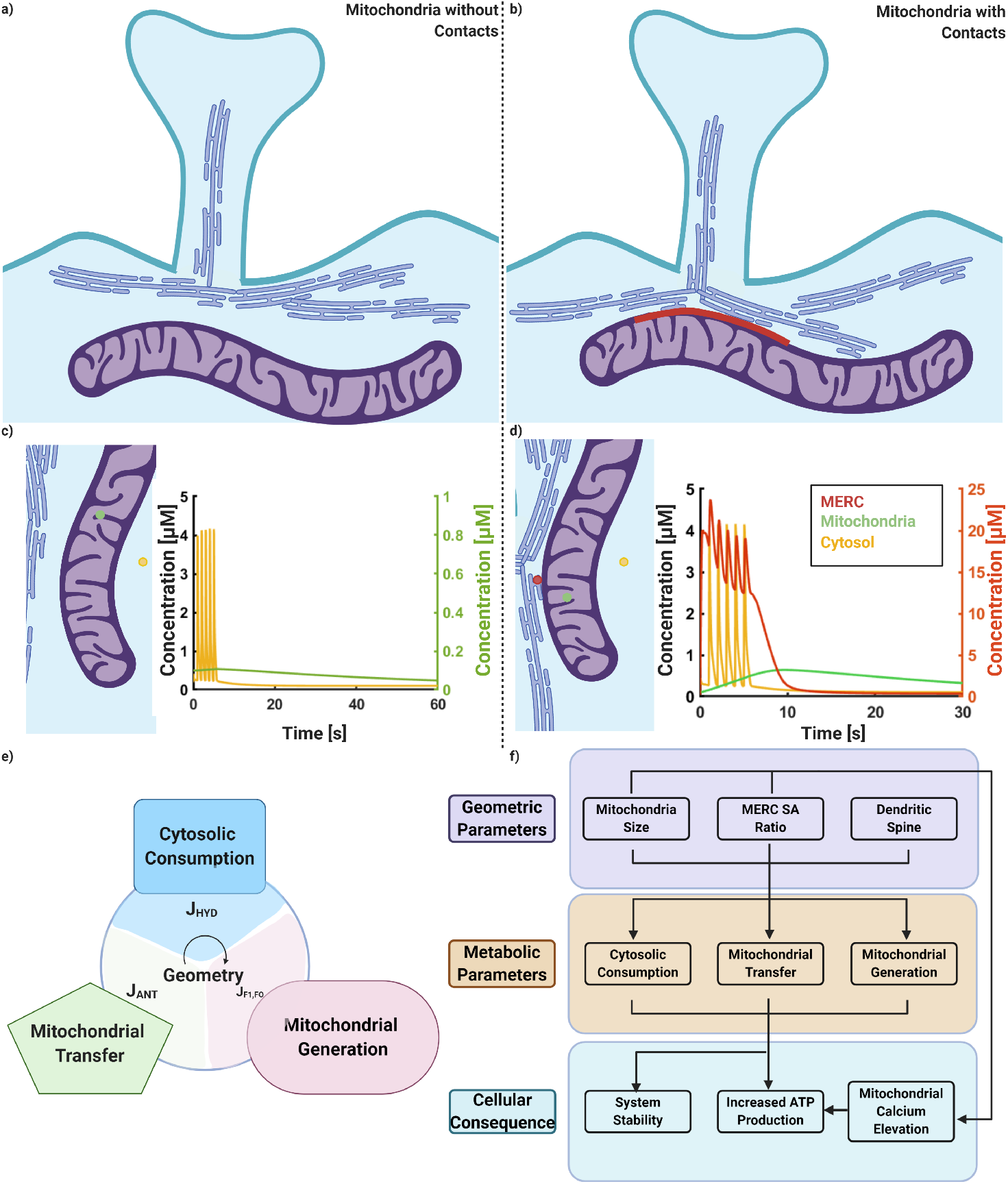
Schematic for the impact of MERCs and geometric regulation of mitochondiral ATP production. **a**) Schematic for dendrite with ER not forming contacts with mitochondria. **b**) Schematic for dendrite with ER forming contacts with mitochondria, highlighted in red **c**) Calcium profiles for cytosol (in yellow) and mitochondria (in green) for the system without any contact sites. **d**) Calcium profiles for cytosol (in yellow), mitochondria (in green), and MERC (in red) for system with contact sites. **e**) Schematic outlining key fluxes that balance ATP in the mitochondria and cytosol explored in Figure 6. **f**) Flow diagram describing the interconnected influence of cell and organelle geometry on metabolic production with added feedback through calcium signaling.

Recently, the mitochondrial dynamics, including fission [47], ROS generation [55], cytochrome c release [56] and MERCs [50] have been implicated in postsynaptic response to different stimuli, particularly in the context of energy requirements of synaptic transmission. Furthermore, the presence of MERCs has been noted as a critical factor in the mitochondrial response and loss of MERCs shows a reduced mitochondrial Ca^2+^ response in neurons [50]. Dendritic mitochondria can range in size from 1 *μ*m to over 10 *μ*m in length (median 2.5 *μ*m) [11, 39, 47] While our spatial model is not to scale (we chose a 1.5 *μ*m long dendrite segment for computational ease), we note that our model predicts that dendritic energy availability is directly proportional to mitochondrial length. Divakaruni et al. report that mitochondrial length is linearly proportional to dendritic length [47]; thus the scaling we find here suggests a common size principle for dendritic energy availability. Separately, Hirayabashi and colleagues [50] reported that PDZD8 is required for MERC formation and that the MERC area fraction ranges from 0.02-0.16 in HeLa cells. We note that increasing MERC to mitochondria surface area ratios has influence over the ATP production dynamics as well as the total ATP production. Although MERCs can have a substantial impact on mitochondria ATP generation in contrast to cases without MERC, the MERC area fraction does not show a linear dependence on ATP production as was observed for mitochondrial size. This implies that the benefit of MERCs is closer to that of a switch-like behavior rather than graded behavior. We also show that biochemical parameters associated with metabolism such as ATP production, nucleotide transport, and ATP hydrolysis also play an important role in cytosolic energy availability by allowing for a graded behavior. Thus, we find that there is a unique balance between energetic and biochemical aspects of mitochondrial function in dendrites. This balance is further regulated in cells by Ca^2+^-dependent mitochondrial fission to complete the feedback loop of signaling and mitochondrial size regulation and will require a deeper understanding of the underlying coupled mechanochemical processes [57].

We note that our model has made some simplifying assumptions to keep the simulations tractable. First, all our geometries are simplified, allowing us to construct a spatial model of multiple organelles. We note that current efforts include using realistic geometries of spines and organelles derived from microscopy to investigate how these play a role in Ca^2+^ and ATP dynamics [52, 58, 59]. Second, we note that we assume that Ca^2+^ and ATP are present in large enough amounts to justify a deterministic approach. We recognize that a stochastic approach would likely give rise to insights in limit cases of few molecules or in noisy environments [37, 38]. Finally, we note that a more complete Ca^2+^ influx model would account for voltage-gated calcium channel dynamics as well [22, 24, 60, 61]. This is a focus of technical development and ongoing effort in our group.

To summarize, we show using a spatial model of Ca^2+^ dynamics in dendrites, including the ER, the mitochondria, and MERCs, that spatial organization of organelles and their contacts plays a critical role in determining the downstream response of a spine to a stimulus. This work sets the stage for future investigation on the coupling between biochemical signaling [62], biophysical mechanisms of organelle transport and tethering, and metabolic pathways, giving a glimpse into the complex regulation of neuronal processes.

## Abbreviations

(ER): Endoplasmic reticulum
(IP_3_): Inositol 1,4,5-trisphosphate
(MERC, also MAM): Mitochondrial Endoplamic Reticulum Contact
(IP_3_R): IP_3_ receptor
(RyR): Ryanodine receptor
(SERCA): sarcoplasmic/endoplasmic reticulum calcium ATPase
(ATP): Adenine triphosphate
(OMM): outer mitochondrial membrane
(IMS): intermembrane space
(MCU): Mitochondrial Calcium Uniporter
(PMCA): Plasma Membrane Calcium ATPase
(IMM): inner mitochondrial membrane
(ETC): Electron Transport Chain
(AGC): Aspartate/glutamate carrier
(MAS): Malate/asparate shuttle
(ANT): Adenine nucleotide translocator
(PDH): Pyruvate dehydrogenase
(GPDH): Glyceraldehyde 3-phosphate dehydrogenase
(NCX): Na^+^/Ca^2+^ exchanger
(mGluR): Metabotropic glutamate receptor
(NMDAR): N-Methyl d-Aspartate Receptor
(PKC): Protein kinase C
(DAG): Diacyl-glycerol
(PSD): Post synaptic density
(PIP2): Phosphatidylinositol (4,5)-bisphosphate

## Author Contributions

A. Leung, D. Ohadi, and P. Rangamani conceived the project. D. Ohadi and A. Leung developed spatial model. A. Leung, G. Pekkurnaz and P. Rangamani conducted simulations and data analysis. All authors contributed to writing the manuscript. A. Leung generated all figures.

## Acknowledgments

The authors would like to acknowledge Justin Laughlin and Miriam Bell for their useful discussion and computational resources. Additionally, we acknowledge biorender.com for tools used to generate figures. Finally, this work was supported by the University of California, San Diego Interfaces Graduate Training Program and a San Diego Fellowship, Air Force Office of Scientific Research (AFOSR) Multidisciplinary University Research Initiative (MURI) grant FA9550-18-10051 (A. Leung, P. Rangamani)

## Supplementary Material

### Mathematical Model Development

In this section, we discuss the details of the model development, assumptions, parameters, and numerical methods.

### Geometry Development

We built a model with five compartments: the postsynaptic density (PSD), the cytosol, one mitochondria, the ER, and a mitochondria ER Contact region (MERC) (Figure 1).

In our simplified geometry, the dendritic spine is modeled as a sphere attached to the dendritic shaft by the spine neck. The spine neck and the dendritic shaft were modeled as cylinders. (Table S1)

Although the morphology of a spine has been shown to govern the magnitude and stability of calcium transients in dendrites in previous studies [24, 31], in this study we simplify complex spine morphology in idealized geometries to focus on the role of mitochondria in neuronal calcium dynamics. While the ER is distributed throughout the cytoplasm, we approximate the ER within the dendritic shaft as long cylinders, keeping volumetric ER to cytoplasm ratios constant. The sizes of these compartments are given in Table S1. While we acknowledge that these simplifications do not capture geometric complexity of a dendrite [48], they allow us to explore the role of spatial organization of the spine in a computational framework.

### Neuronal Stimulus Patterns

While there are a multitude of signaling frequencies and patterns, in this work we focus on a simple periodic stimulus, 1 Hz for 5 seconds [63] to emulate signaling in an established spine. Mathematically, we approximate glutamate stimulus as discrete pulses in a well-mixed system within COMSOL. Since we are not modeling synaptic vesicles or presynaptic neuron signaling, we apply a series of delta functions temporally spaced according to the desired pulse train (i.e. 1 Hz stimulus has delta functions spaced 1 seconds apart). Glutamate then decays with a time constant of 1.2ms [64]. Key contributing factors to this decay are rapid reabsorbed by the signaling neurons and astrocytic cells as well as diffusion [65].

### Calcium Dynamics

After glutamate binds to receptors on the post-synaptic density (PSD), there is a large calcium influx from the extracellular space and endoplasmic reticulum. This signaling initiates at the dendritic spine and propagates to the dendritic shaft, where the mitochondria are primarily located. We assume that the extracellular calcium does not deplete and does not alter the flux of synaptic calcium signaling. Extracellular calcium concentrations are typically several orders of magnitude higher than intracellular calcium [66].

The calcium dynamics in our model are based on work done by [25, 27, 67], and parameters were adapted and suitably modified to reflect neuronal calcium metabolism.

- **Postsynaptic area and Receptor Dynamics:** This compartment includes membrane-bound molecules on the postsynaptic area: NMDA receptor (NMDAR), free mGluRs (R_2_), glutamate-bound mGluRs (DIM), phosphorylated mGluRs (DIM_*p*_), and second messenger DAG. These molecules localize to the PSD area of spine heads. Calcium enters the spine head through NMDAR. The NMDAR dynamics are described in a multistate model [68]. And the NMDAR-induced calcium influx is modeled as:

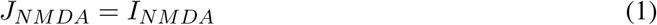

where, *I*_*NMDA*_ is the current associated with calcium influx. Mathematical descriptions of all terms are expanded in Table S2. Glutamate binds to the mGluR on the spine head, which induces G_*qα*_ to separate from G_*qβγ*_ and activate PLC_*β*_ enzyme. PLC_*β*_ enzyme cleaves PIP_2_ (a membrane phospholipid) and generates DAG and IP_3_. While DAG remains on the membrane and activates PKC, which then may phosphorylates mGluR, phosphotases dephosphorylate mGluR. This phosphorylation can inhibit the dimerization of mGluR IP_3_ diffuses through the cytosol and initiates Ca^2+^ release from internal stores by binding to receptors (IP_3_R) on the ER membrane. IP_3_ in the cytosol is degraded to IP_2_ and IP_4_. The degradation is assumed to recycle phospholipids such that IP_3_ production is not rate limited by PIP_2_ availability. Although not modeled here, Ca^2+^ can also be released from ER by Ryanodine receptors and conversely can be actively transported back into the ER by Sarcoplasmic/endoplasmic reticulum calcium-ATPase (SERCA) on the ER membrane. Equations associated with the mGluR cascade are included in Table S2. The calcium flux is modeled as a boundary condition on the endoplasmic reticulum membrane, in contrast to the NMDAR on the PSD.
- **Cytosolic Calcium Dynamics:** Ca^2+^ transients that accumulate due to influx at the PSD are able to diffuse throughout the cytosol. While we do not explicitly model calcium buffers, buffering capacity is included through boundary conditions between compartments. Cytosolic Ca^2+^ dynamics in 3D is defined as:

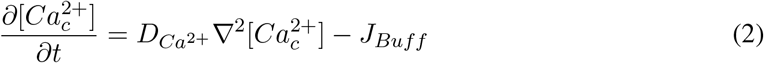

where *D*_*Ca*^2+^_ is diffusion coefficient and ∇^2^ is Laplacian operator in 3D. The boundary condition for Ca^2+^ influx through NMDAR in postsynaptic area and Ca^2+^ efflux from cytosol is defined as,

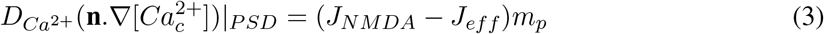

where *m*_*p*_ is a geometric factor for fluxes on the boundary and is defined as the ratio of the cytosol volume to the postsynaptic surface area. J_*NMDA*_ is defined as the flux as a result of the NMDA receptor activity. J_*eff*_ is defined as the activity as a result of plasma membrane calcium ATP-ase. The boundary fluxes on ER for cytosolic Ca^2+^ are defined as:

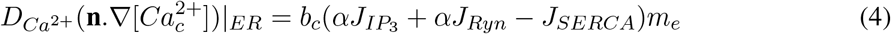

in which b_*c*_ is the buffering capacity of cytosol and *α* is ER to cytosol volume ratio. *J*_*IP R*_ is the Ca^2+^ flux from the ER into the cytosol through the IP_3_ receptor, *J_RY R_* is the Ca^2+^ flux from the ER into the cytosol through the Ryanodine receptor, *J*_*SERCA*_ is the Ca^2+^ flux from the cytosol into the ER through the SERCA ATPase pumps, and *m*_*e*_ is a geometric factor for fluxes on the boundary and is defined as the ratio of the cytosolic volume to the ER surface area. *J*_*IP_3_R*_ and *J*_*RY R*_ are defined with respect to ER volume and *J*_*SERCA*_ is defined with respect to the cytosol volume. The mitochondrial boundary fluxes for cytosolic Ca^2+^ are defined as:

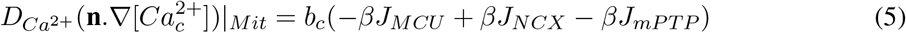

in which *β* is mitochondria to cytosol volume ratio. *J*_*MCU*_ is Ca^2+^ flux from the cytosol into the mitochondrion through the MCU channel, *J*_*NCX*_ is Ca^2+^ flux from the mitochondrion into the cytosol through Na^+^/Ca^2+^ exchanger, and *J*_*mPTP*_ is Ca^2+^ bidirectional flux of the mitochondrial Permeability Transition Pore. All of the fluxes within Eq. 5 are defined with respect to mitochondrion volume.
- **Calcium buffering:** Calcium buffering is included both explicitly and implicitly. The explicit calcium buffers as diffusive buffering molecules and immobile buffers are included as boundary conditions at the plasma membrane.

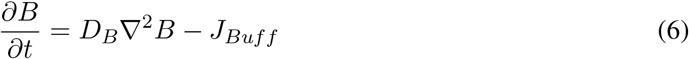

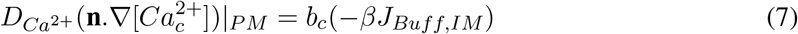 Implicitly, we consider buffering terms within the boundary conditions between compartments. These buffering terms are denoted as *b*_*m*_, *b*_*c*_, *b*_*er*_ and *b*_*MERC*_.
- **Mitochondrial Calcium Dynamics:** The spatiotemporal dynamics of mitochondrial Ca^2+^ is given by

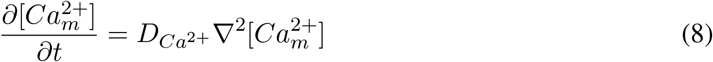 The boundary fluxes on mitochondrion surface are defined as:

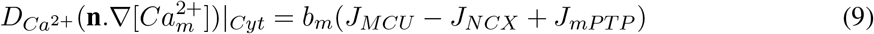

in which *b*_*m*_ is the buffering capacity of Ca^2+^ in the mitochondrion. We do not consider the distinction between the outer and inner mitochondrial membrane. Voltage-dependent anion channels transport calcium highly efficiently during neuronal activation. Thus, we may assume, as done in [26] [25], that the calcium concentrations in the cytosol and outer mitochondrial calcium approach a rapid equilibrium. The inner mitochondrial membrane calcium fluxes are dependent solely on the cytosolic calcium. The same applies to fluxes regarding the mitochondria ER contact microdomain, which is represented in this model as a separate compartment.
- **ER Calcium Dynamics:** The only variable in ER compartment is 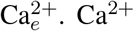 in the ER is defined as:

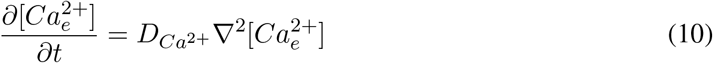 The boundary fluxes on ER are defined as:

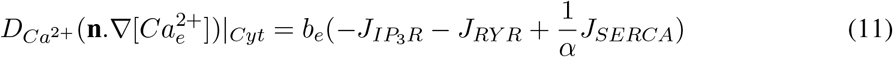

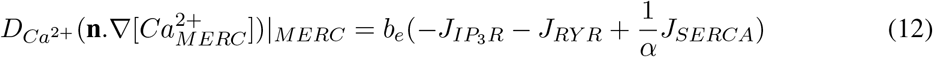

in which *b*_*e*_ is the buffering capacity of ER. While the endoplasmic reticulum is a sprawling network of interconnected membranes containing a wide range of proteins and ribosomes, we simplify this network into a series of tubes that span the length of the model dendritic shaft.
- **Mito-ER Contacts:** The ER is known to closely contact the mitochondria and form junctions known as mitocondria-ER Contacts (henceforth referred to as MERC) [17] [46]. This close proximity, which can be as small as 10 nm and is filled with actin [69],calcium binding proteins [18], and pathways that inhibit calcium reuptake [46]. Therefore we model the calcium dynamics in these regions as a separate microdomain from the cytosol. In order to activate IP_3_R on the ER surface facing this microdomain, IP_3_ must also be present. Thus there are 2 main components within this microdomain, Ca^2+^ and IP_3_. We do not model the molecular composition of the MERC microdomain that control the influx and efflux of Ca^2+^. Rather, we use the diffusive fluxes at the MERC boundaries as lumped parameters to reflect the different protein distributions across the MERC boundaries. We model the geometry of this microdomain as a shell that surrounds the circumference of the mitochondria and connects to the endoplasmic reticulum. This shape was chosen to reflect EM images of MERCs and to efficiently model the portion of ER in contact with the mitochondria. The calcium concentration within this compartment is given by:

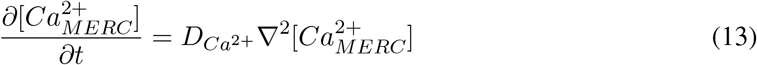 And the boundary conditions for the interface between the MERC and the ER, Mitochondria, and cytosol are given as follows:

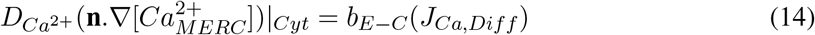

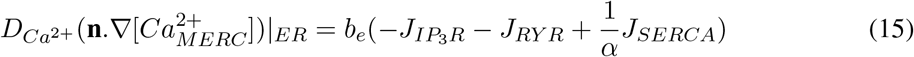

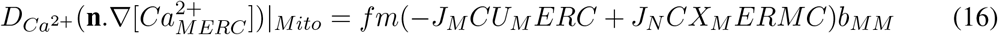

### Energetic considerations in the model

In this model, we focus on the impact on calcium on mitochondrial ATP production, as done in [25]. Our metabolic model focuses on NAD and NADH within the mitochondria, and ATP and ADP in both the mitochondria and cytosol. To avoid modeling the complex spatial patterns of the inner mitochondrial membrane, we assume that ADP conversion into ATP is a volumetric reaction within the mitochondria. Redox and mitochondrial potential are likewise modeled as volumetric terms within the mitochondria.

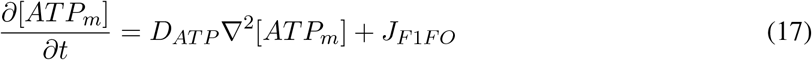

ATP and ADP are exchanged through the Adenine Nucleotide Transporter (ANT), which is modeled as a boundary condition.

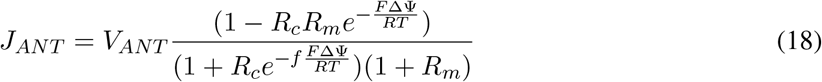

After transfer into the cytosol, ATP freely diffuses and is hydrolyzed by unspecified reactions in the cytosol.

### Numerical Methods

The Ca^2+^ concentrations and other variables concentrations in all compartments are high enough to be modeled through a deterministic approach. Simulations were conducted using the commercially available finite-element software COMSOL Multiphysics 5.4 [70]. In order to solve our system of partial differential equations, we used time-dependent general partial differential equations and general boundary partial differential equations modules [70]. Starting with a coarse and unstructured mesh, we decreased the mesh size until we obtained the same results when using the maximum mesh size. COMSOL was allowed to optimize the element sizes through the “physics-controlled mesh” option. The linear system was solved directly by using the PARADISO solver on a Linux-based compute cluster. Newton’s method (nonlinear method) was used to linearize the system. Time integration was performed using a backward differentiation formula (BDF) with both adaptive order and adaptive step sizes. All COMSOL source files will be available on Rangamani lab Github page upon publication.

## Supplementary tables

**Table S1:** Model geometry

| Organelle | Shape | Length ($\mu\text{m}$) | Diameter ($\mu\text{m}$) | Surface area ($\mu\text{m}^2$) | Volume ($\mu\text{m}^3$) |
| --- | --- | --- | --- | --- | --- |
| Dendrite | cylinder | 1.6 | 0.6 |  |  |
| Mitochondrion | cylinder | 0.6 | 0.3 |  |  |
| Tubule mitochondrion | cylinder | 1.2 | 0.21 |  |  |
| Each of 4 tubule ER | cylinder | 1.5 | 0.08 |  |  |
| Spine 1 (head) | sphere | - | 0.6 |  | 0.058 [71] |
| Spine 1 (neck) | cylinder | 0.4 | 0.2 |  |  |
| Spine 1 (PSD) | spherical cap |  |  | 0.132 [72] |  |
| Spine 2 (head) | sphere | - | 0.4 |  | 0.028 [71] |
| Spine 2 (neck) | cylinder | 0.5 | 0.1 |  |  |
| Spine 2 (PSD) | spherical cap |  |  | 0.055 [72] |  |
| Spine apparatus (head) | sphere | - | 0.34 |  |  |
| Spine apparatus (neck) | cylinder | 0.5 | 0.06 |  |  |
| Spine apparatus (base) | cylinder | 1.5 | 0.08 |  |  |
| Distance between spine 1 and spine 2 | - | 1 | - |  |  |
| Distance between the ER and mitochondria | - | 0.050 [41] | - |  |  |
|  | - | 0.030 [73] | - |  |  |

**Table S2:**
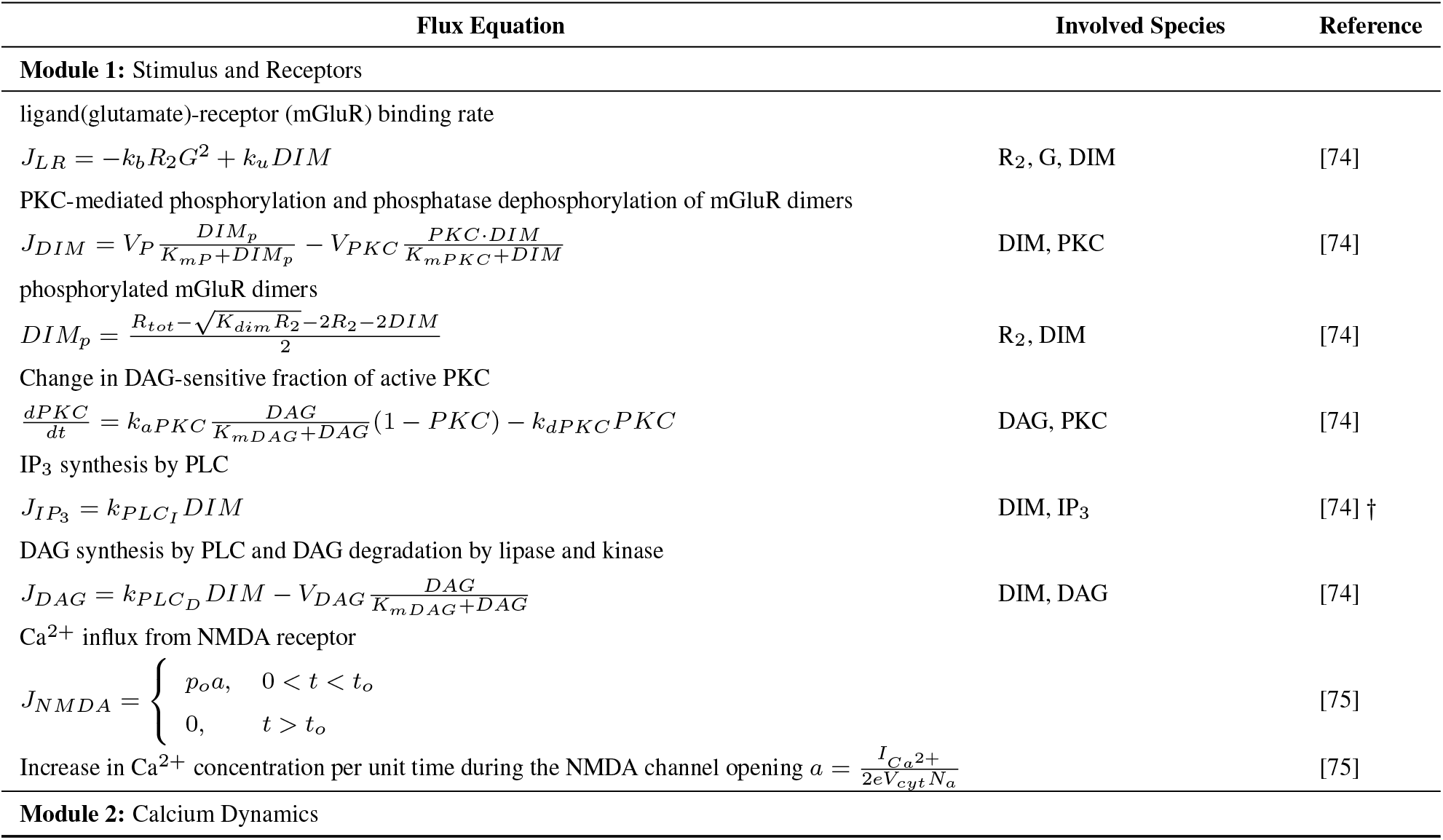
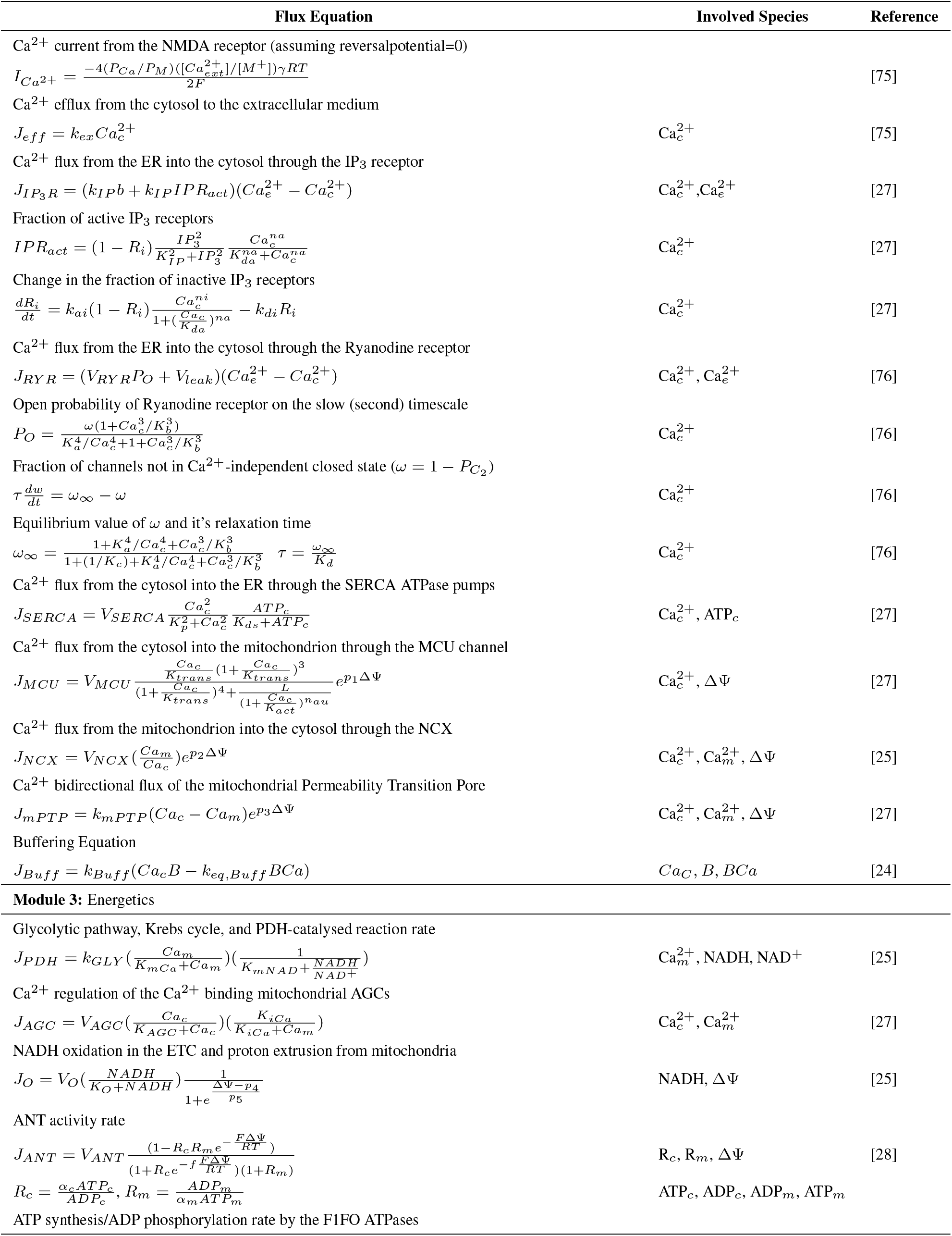
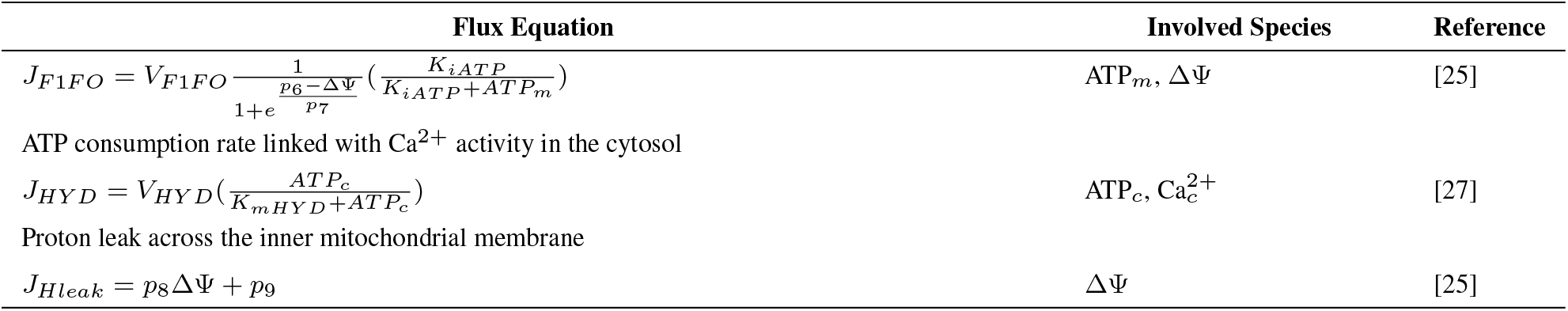
Fluxes and Description

**Table S3:**
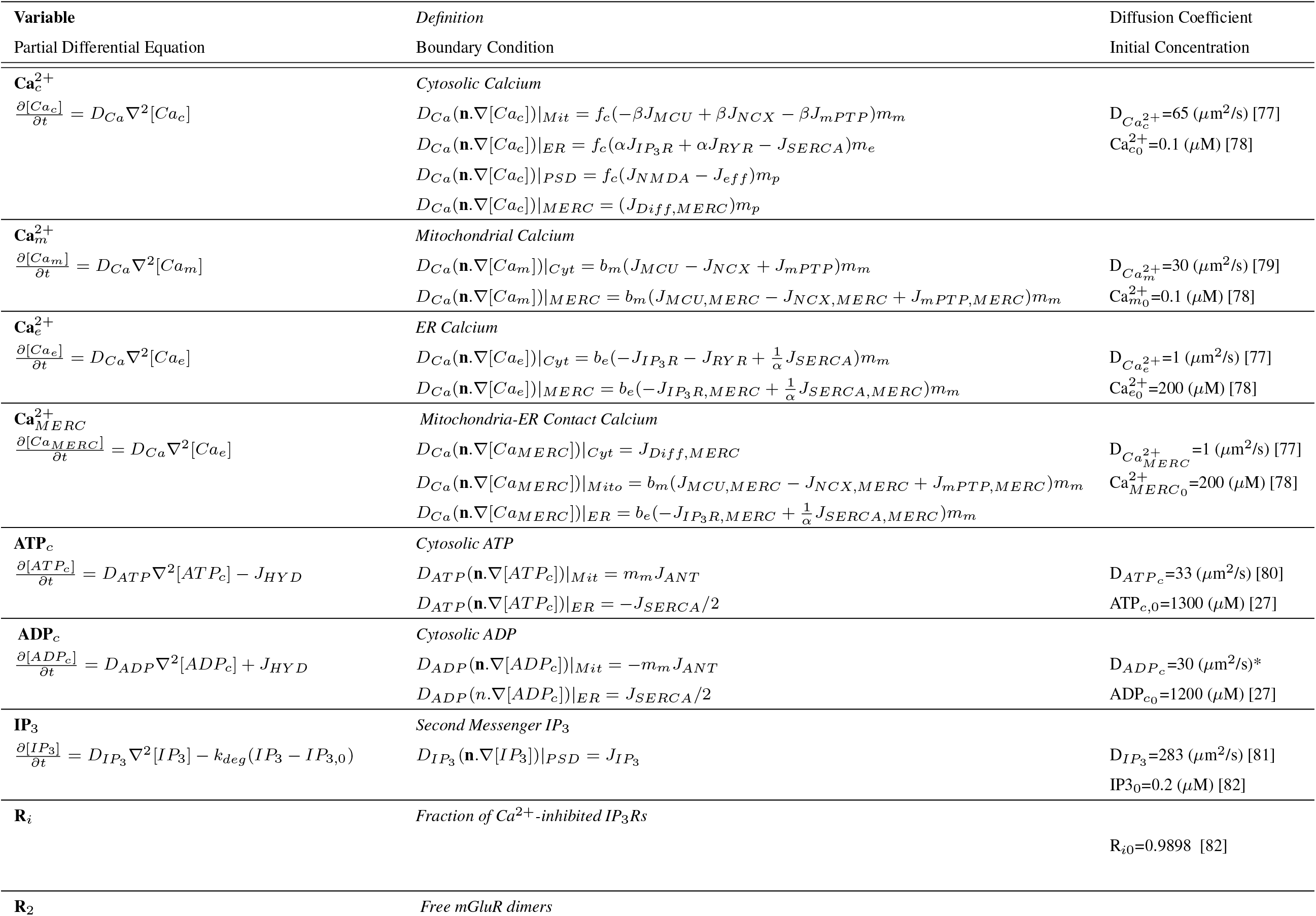
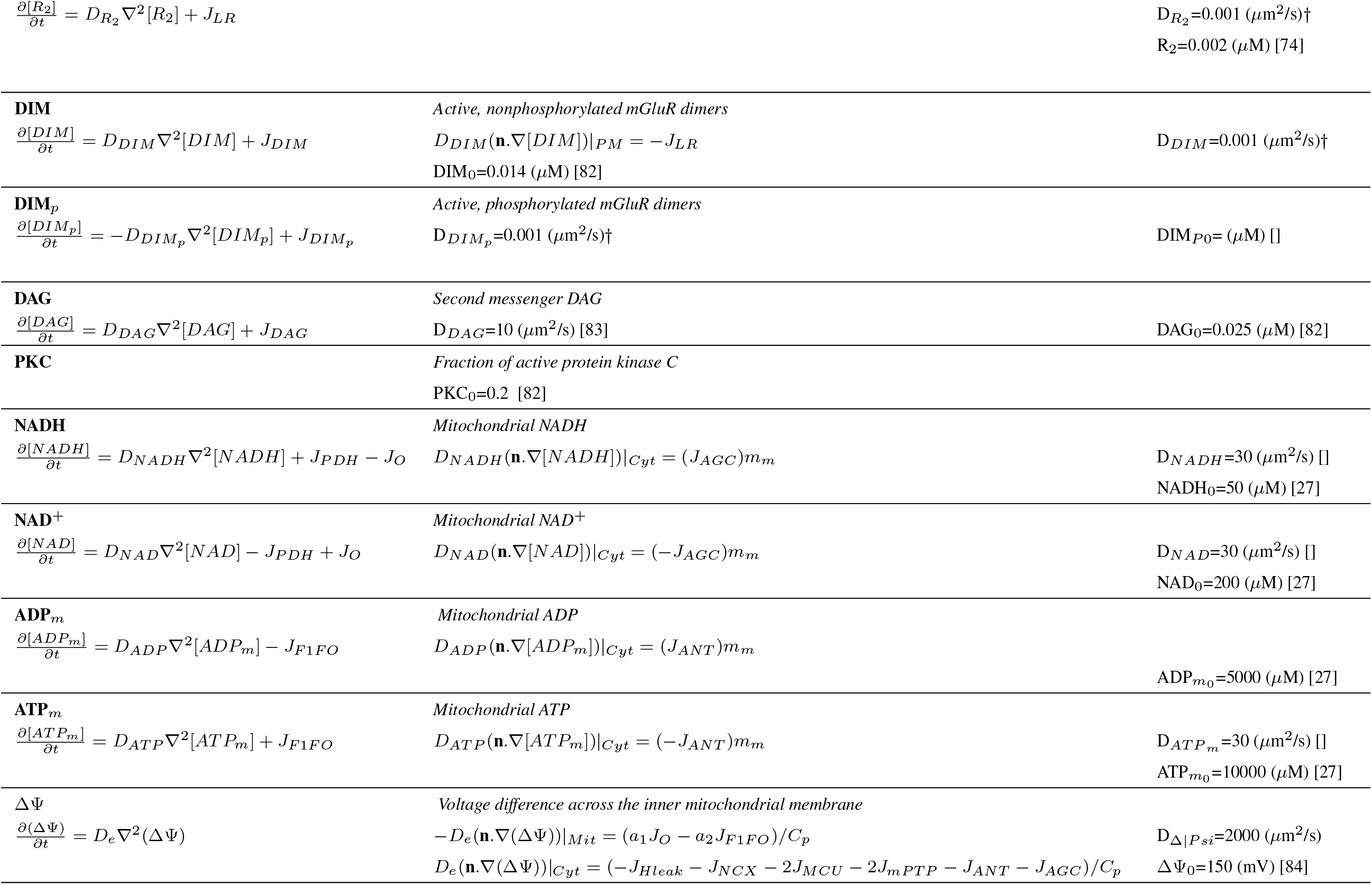
Initial conditions, boundary conditions and partial differential equations

**Table S4:** Peak Times for Figure 4

| Mitochondrial Length [$\mu\text{m}$] | $\text{ATP}_M$ Peak Time [s] | $\text{ATP}_C$ Peak Time [s] |
| --- | --- | --- |
| 0.1 | 6.1 | 0 |
| 0.2 | 10.2 | 0 |
| 0.4 | 11.7 | 0 |
| 0.6 | 12.0 | 0 |
| 0.8 | 12.3 | 22.4 |
| 1.0 | 12.7 | 32.1 |
| 1.2 | 13.6 | 39.6 |

**Table S5:** Model Parameters

| Flux | Parameter | Definition | Value | Units | Reference |
| --- | --- | --- | --- | --- | --- |
| | $b_c$ | buffering capacity of cytosol | 0.01 | - | |
| | $b_m$ | buffering capacity of mitochondria | 0.01 | - | |
| | $b_e$ | buffering capacity of ER | 0.0003 | - | |
| | $m_p$ | cytosol volume to postsynaptic surface area ratio | * | - | |
| | $m_m$ | cytosol volume to mitochondrion surface area ratio | * | - | |
| | $m_e$ | cytosol volume to ER surface area ratio | * | - | |
| | $\alpha$ | ER volume to cytosol volume ratio | * | - | |
| | $\beta$ | mitochondria volume to cytosol volume ratio | * | - | |
| | $\kappa_i$ | endogenous buffer capacity | 20 | C | [75] |
| $J_{NMDA}$ | $p_o$ | open probability of NMDAR | 0.3 | - | [75] |
| $J_{NMDA}$ | $t_o$ | open time of NMDAR | 100 | ms | [75] |
| $J_{NMDA}$ | $e$ | elementary charge | $1.6 \times 10^{-19}$ | - | [75] |
| $J_{NMDA}$ | $N_a$ | Avogadro number | $6.022 \times 10^{23}$ | $\text{mol}^{-1}$ | [75] |
| $J_{NMDA}$ | $V_{cyt}$ | cytosolic volume | | | [75] |
| $J_{NMDA}$ | $P_{Ca}/P_M$ | permeability of monovalent ions | 3.6 | | [75] |
| $J_{NMDA}$ | $[\text{Ca}_{ext}^{2+}]$ | concentration of extracellular $\text{Ca}^{2+}$ | 2 | mM | [75] |
| $J_{NMDA}$ | $[\text{M}^+]$ | concentration of monovalent ions | 130 | mM | [75] |
| $J_{NMDA}$ | $\gamma$ | Conductance of monovalent ions | 46 | pS | [75] |
| $J_{NMDA}$ | $R$ | gas constant | 8.315 | $\text{JK}^{-1}\text{mol}^{-1}$ | [75] |
| $J_{NMDA}$ | $T$ | absolute temperature | 303 | K | [75] |
| $J_{eff}$ | $k_{ex}$ | $\text{Ca}^{2+}$ clearance rate constant in the absence of $\text{Ca}^{2+}$ buffer | 0.0016 | $\text{s}^{-1}$ | [75] |
| $J_{LR}$ | $k_u$ | kinetic constant of glutamate unbinding to mGluR dimers | 2 | $\text{s}^{-1}$ | [74] |
| $J_{LR}$ | $k_b$ | kinetic constant of glutamate binding to mGluR dimers | 1 | $\mu\text{M}^{-2}\text{s}^{-1}$ | [74] |
| $J_{LR}$ | $G$ | glutamate concentration | 4 | $\mu\text{M}$ | [85] |
| $J_{DIM}$ | $V_P$ | maximal rate of mGluR dephosphorylation by a phosphatase | 0.05 | $\mu\text{Ms}^{-1}$ | [74] |
| $J_{DIM}$ | $K_{mP}$ | Michaelis-Menten constant of mGluR dephosphorylation by a phosphatase | $5 \times 10^{-4}$ | $\mu\text{M}$ | [74] |
| $J_{DIM}$ | $V_{PKC}$ | maximal rate of mGluR phosphorylation by PKC | 0.2 | $\mu\text{Ms}^{-1}$ | [74] |

Table S5 – continued from previous page
| Flux | Parameter | Definition | Value | Units | Reference |
| --- | --- | --- | --- | --- | --- |
| $J_{DIM}$ | $K_{mPKC}$ | Michaelis-Menten constant of mGluR phosphorylation by PKC | $5 \times 10^{-4}$ | $\mu\text{M}$ | [74] |
| $J_{DIM}$ | $K_{dim}$ | dimerization constant of mGluR | 0.1 | $\mu\text{M}$ | [74] |
| $J_{DIM}$ | $R_{tot}$ | total concentration of mGluR in monomeric form | 0.075 | $\mu\text{M}$ | [74] |
| $J_{DIM}$ | $k_{aPKC}$ | Normalized (divided by PKC total conc.) maximal rate of PKC activation | 0.2 | $\text{s}^{-1}$ | [74] |
| $J_{DIM}$ | $k_{dPKC}$ | rate constant of PKC deactivation | 0.2 | $\text{s}^{-1}$ | [74] |
| $J_{DIM}$ | $K_{mDAG}$ | activation constant of PKC by DAG | 0.06 | $\mu\text{M}$ | [74] |
| $J_{IP_3}$ | $k_{PLC_I}$ | catalytic activity of PLC for $\text{IP}_3$ | 1.25 | $\text{s}^{-1}$ | [74] |
| $J_{IP_3}$ | $k_{DAG_I}$ | DAG enhancing $\text{IP}_3$ production via its positive feedback effect on PLC activity | 8 | - | † |
| $J_{IP_3}$ | $k_{dI}$ | rate constant of $\text{IP}_3$ degradation (to $\text{IP}_2$ and $\text{IP}_4$) | 0.12 | $\text{s}^{-1}$ | [74] |
| $J_{DAG}$ | $k_{PLC_D}$ | catalytic activity of PLC for DAG | 1.25 | $\text{s}^{-1}$ | [74] |
| $J_{DAG}$ | $V_{DAG}$ | maximal rate of DAG degradation | 0.0325 | $\mu\text{Ms}^{-1}$ | [74] |
| $J_{DAG}$ | $K_{DAG}$ | Michaelis-Menten constant of DAG degradation | 0.012 | $\mu\text{M}$ | [74] |
| $J_{R_YR}$ | $V_{R_YR}$ | rate constant of the $\text{Ca}^{2+}$ flux from the ER into the cytosol through the RYR | 5 | $\text{s}^{-1}$ | [76] |
| $J_{R_YR}$ | $V_{leak}$ | basal flux in the absence of RYR receptors/ rate of $\text{Ca}^{2+}$ leak from the ER? | 0.15 | $\text{s}^{-1}$ | [76] |
| $J_{R_YR}$ | $K_a^4$ | RYR dissociation constant | 0.019 | $\mu\text{M}^4$ | [76] |
| $J_{R_YR}$ | $K_b^3$ | RYR dissociation constant | 0.257 | $\mu\text{M}^3$ | [76] |
| $J_{R_YR}$ | $K_c$ | RYR dissociation constant | 0.057 | - | [76] |
| $J_{R_YR}$ | $K_d$ | rate constant for transition between open and closed state | 0.1 | $\text{s}^{-1}$ | [76] |
| $J_{IP_3R}$ | $k_{IP}$ | rate constant of the $\text{Ca}^{2+}$ flux from the ER into the cytosol through the $\text{IP}_3\text{R}$ | 30 | $\text{s}^{-1}$ | [27] |
| $J_{IP_3R}$ | $k_{IPb}$ | basal flux in the absence of $\text{IP}_3$ receptors/ rate of $\text{Ca}^{2+}$ leak from the ER | 0.3 | $\text{s}^{-1}$ | [27] |
| $J_{IP_3R}$ | $K_{IP}$ | dissociation constant of $\text{IP}_3$ binding to $\text{IP}_3$ receptor | 1 | $\mu\text{M}$ | [27] |
| $J_{IP_3R}$ | $K_{da}$ | dissociation constant of $\text{Ca}^{2+}$ binding to the activating site of $\text{IP}_3$ receptor | 0.3 | $\mu\text{M}$ | [27] |
| $J_{IP_3R}$ | $n_a$ | Hill coefficient for cooperative $\text{Ca}^{2+}$-binding of $\text{IP}_3$ receptor activation | 3 | - | [27] |
| $J_{IP_3R}$ | $n_i$ | Hill coefficient for cooperative $\text{Ca}^{2+}$-binding of $\text{IP}_3$ receptor inhibition | 4 | - | [27] |
| $J_{IP_3R}$ | $k_{ai}$ | association constant of $\text{Ca}^{2+}$-binding to the inhibitory site of $\text{IP}_3$ receptor | 20 | $\mu\text{M}^4.\text{s}^{-1}$ | [27] |
| $J_{IP_3R}$ | $k_{di}$ | dissociation constant of $\text{Ca}^{2+}$-binding to the inhibitory site of $\text{IP}_3$ receptor | 0.02 | $\text{s}^{-1}$ | [27] |
| $J_{SERCA}$ | $V_{SERCA}$ | maximal rate of the SERCA pumps | 120 | $\mu\text{M}.\text{s}^{-1}$ | [27] |
| $J_{SERCA}$ | $K_p$ | $\text{Ca}^{2+}$ dissociation constant from the SERCA pumps | 0.35 | $\mu\text{M}$ | [27] |
| $J_{SERCA}$ | $K_{ds}$ | ATP dissociation constant from the SERCA pumps | 0.05 | $\mu\text{M}$ | [27] |
| $J_{MCU}$ | $V_{MCU}$ | maximal rate of the mitochondrial $\text{Ca}^{2+}$ uniporter | 0.006 | $\mu\text{M}.\text{s}^{-1}$ | [86] |
| $J_{MCU}$ | $K_{trans}$ | dissociation constant of $\text{Ca}^{2+}$ transport in uniporter | 6 | $\mu\text{M}$ | [86] |
| $J_{MCU}$ | $K_{act}$ | dissociation constant of uniporter activation by $\text{Ca}^{2+}$ | 0.38 | $\mu\text{M}$ | [86] |
| $J_{MCU}$ | $L$ | allosteric constant for transition between relaxed and taut form in four equivalent binding domains of the uniporter | 50 | - | [86] |
| $J_{MCU}$ | $n_{au}$ | uniporter activation cooperativity parameter | 2.3 | - | [27] |
| $J_{MCU}$ | $p_1$ | uniporter voltage dependence parameter | 0.1 | $\text{mV}^{-1}$ | [27] |
| $J_{NCX}$ | $V_{NCX}$ | maximal rate of the NCX | 0.35 | $\mu\text{M}^{-1}$ | [27] |
| $J_{NCX}$ | $p_2$ | exchanger voltage dependence parameter | 0.016 | $\text{mV}^{-1}$ | [27] |
| $J_{mPTP}$ | $k_{PTP}$ | rate constant of the bidirectional $\text{Ca}^{2+}$ flux through the mPTP | 0.008 | $\text{s}^{-1}$ | [27] |
| $J_{mPTP}$ | $p_3$ | bidirectional $\text{Ca}^{2+}$ leak voltage dependence parameter | 0.05 | $\text{mV}^{-1}$ | [27] |
| $J_{PDH}$ | $k_{GLY}$ | GPDH reaction rate which reflects the flux through the glycolytic pathway | 450 check | $\mu\text{M}.\text{s}^{-1}$ | [27] |
| $J_{PDH}$ | $K_{mCa}$ | Michaelis-Menten constant for Krebs cycle activation by mitochondrial $\text{Ca}^{2+}$ | 0.1 | $\mu\text{M}$ | [27] |

Table S5 – continued from previous page
| Flux | Parameter | Definition | Value | Units | Reference |
| --- | --- | --- | --- | --- | --- |
| $J_{PDH}$ | $K_{mNADH}$ | Michaelis-Menten constant for $NAD^+$ consumption in the Krebs cycle | 1 | $\mu M$ | [27] |
| $J_{AGC}$ | $V_{AGC}$ | maximal rate of NADH production via MAS | 25 | $\mu M.s^{-1}$ | [27] |
| $J_{AGC}$ | $K_{AGC}$ | cytosolic $Ca^{2+}$ dissociation constant from the AGC | 0.14 | $\mu M$ | [27] |
| $J_{AGC}$ | $K_{iCa}$ | inhibition constant of AGS by cytosolic $Ca^{2+}$ | 0.3 | $\mu M$ | [86, 87] |
| $J_O$ | $V_O$ | maximal rate of NADH oxidation in the ETC | 600 | $\mu M.s^{-1}$ | [27] |
| $J_O$ | $K_O$ | Michaelis-Menten constant for NADH oxidation in the ETC | 100 | $\mu M$ | [27] |
| $J_O$ | $p_4$ | ETC voltage dependence parameter 1 | 177 | mV | [27] |
| $J_O$ | $p_5$ | ETC voltage dependence parameter 2 | 5 | mV | [27] |
| $J_{ANT}$ | $V_{ANT}$ | maximal rate for $ADP_c/ADP_m$ exchange in the absence of ATP | 5000 | $\mu M.s^{-1}$ | [27] |
| $J_{ANT}$ | $\alpha_c$ | cytosolic ADP and ATP buffering parameter | 0.111 | - | [27] |
| $J_{ANT}$ | $\alpha_m$ | mitochondrial ADP and ATP buffering parameter | 0.139 | - | [27] |
| $J_{ANT}$ | $f$ | fraction of $\Delta\Psi$ responsible for the large increase in the Michaelis-Menten constant for ATP uptake by energized mitochondria | 0.5 | - | [27] |
| $J_{F1FO}$ | $V_{F1FO}$ | maximal rate of F1FO ATPase | 35000 | $\mu M.s^{-1}$ | [27] |
| $J_{F1FO}$ | $p_6$ | F1FO ATPase voltage dependence parameter 1 | 190 | mV | [27] |
| $J_{F1FO}$ | $p_7$ | F1FO ATPase voltage dependence parameter 2 | 8.5 | mV | [27] |
| $J_{F1FO}$ | $K_{iATP}$ | F1FO ATPase inhibition constant by mitochondrial ATP | 1000 | $\mu M$ | [27] |
| $J_{HYD}$ | $V_{HYD}$ | maximal rate of ATP hydrolysis | 100 | $\mu M.s^{-1}$ | [27] |
| $J_{HYD}$ | $K_{mHYD}$ | Michaelis-Menten constant for ATP hydrolysis | 1000 | $\mu M$ | [27] |
| $J_{Hleak}$ | $p_8$ | proton leak voltage dependence parameter | 2 | $\mu M.s^{-1}.mV^{-1}$ | [27] |
| $J_{Hleak}$ | $p_9$ | proton leak pH dependence constant | -30 | $\mu M.s^{-1}$ | [27] |

## Supplementary Figures

**Figure S1:**
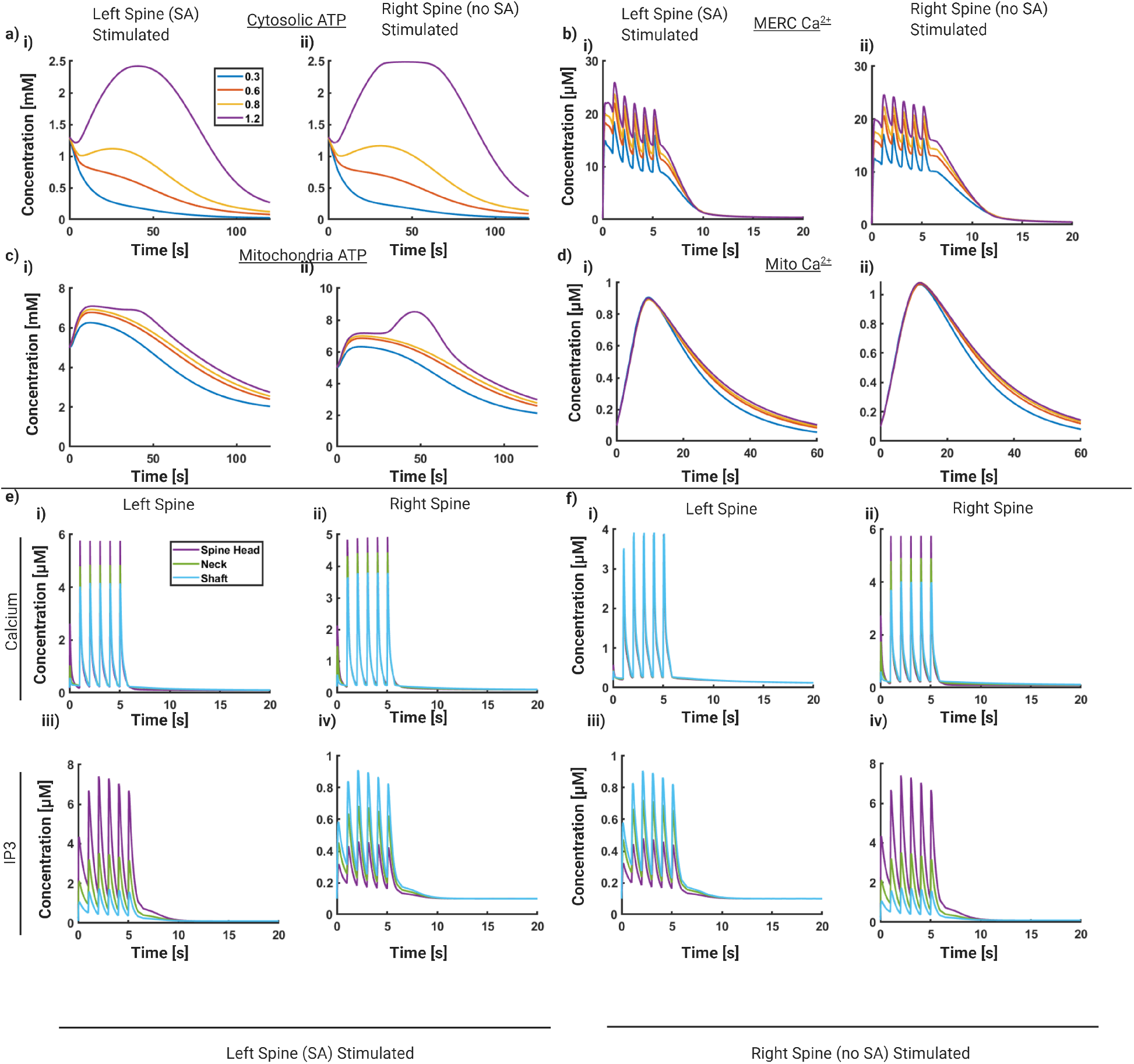
Comparison between stimulus site for larger spine with apparatus and smaller spine without apparatus. **a**) For 4 different mitochondrial sizes, **i)** shows the volume averaged cytosolic ATP concentration when the spine with spine apparatus is stimulated (left spine) and **ii)** shows the average cytosolic ATP concentration when the spine without spine apparatus is stimulated (right spine) **b**) Same as **a**) for volume averaged MERC calcium concentration **c**) Same as **a**) for volume averaged mitochondrial ATP concentration **d**) Same as **a**) for volume averaged mitochondrial calcium **e**) For a single mitochondria size (0.6 micron), we show the stimulated spine and neighboring spine dynamics when the left spine (with SA) is stimulated for: **i)** calcium concentrations for 3 points in left spine head cytosol, **ii)** IP3 concentrations for 3 points in the left spine head cytosol, **iii)** calcium concentrations for 3 points in the right spine head cytosol, and **iv)** IP_3_ concentrations for 3 points in the right spine head cytosol. **f**) Same as **e**) for when the spine on the right with no spine apparatus is stimulated.

**Figure S2:**
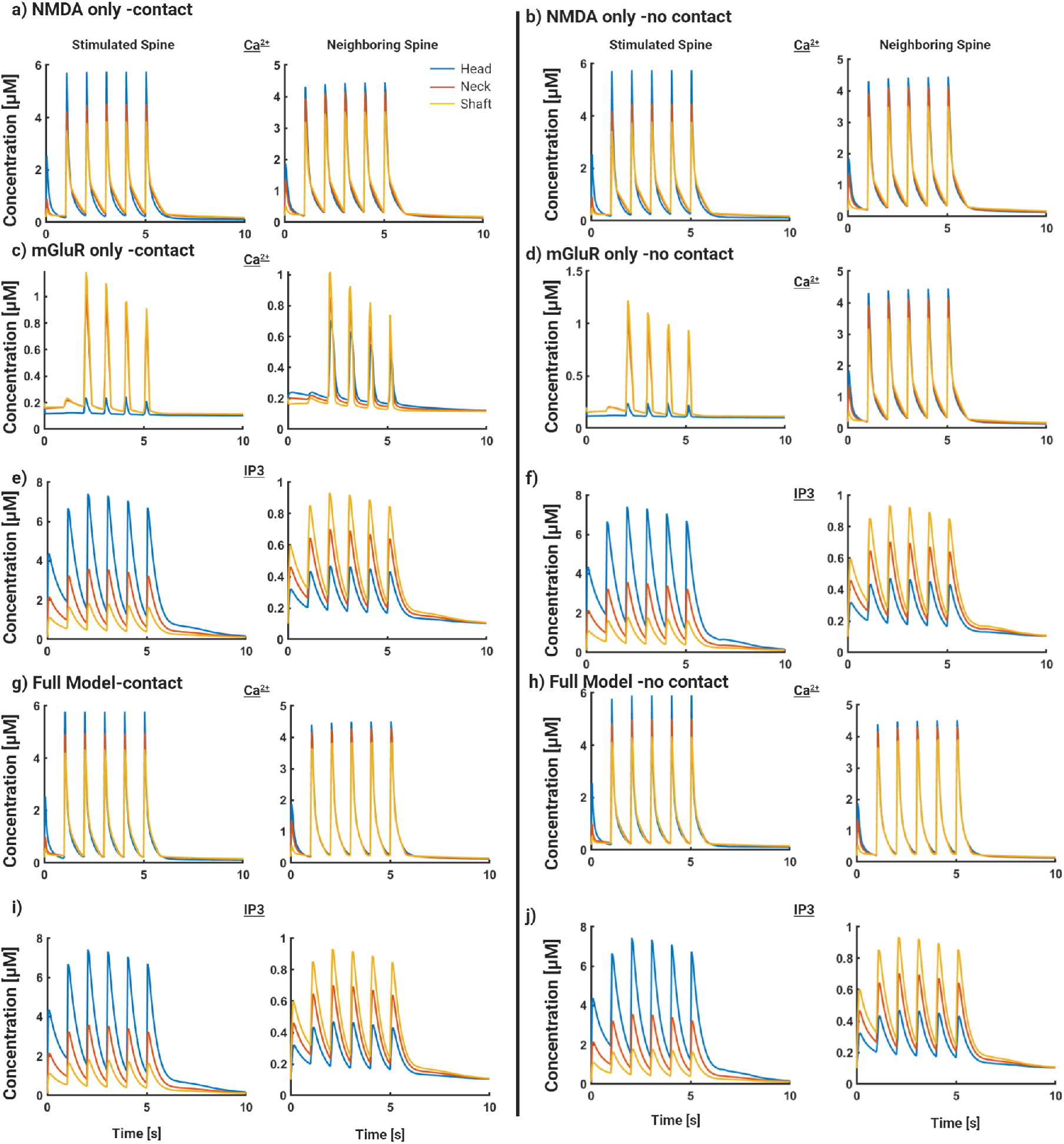
Ca^2+^ and IP_3_ Dynamics during stimulus from mGluR, NMDAR, and both models. **a**) Calcium dynamics taken at 3 points in both the stimulated and neighboring spine in NMDAR-only system with MERC. **b**) same as **a**) for no MERC. **c**) Calcium dynamics taken at 3 points in both the stimulated and neighboring spine in mGluR-only system with MERC. **d**) Same as **c**) with no MERC. **e**)IP_3_ dynamics taken at 3 points in both stimulated and neighboring spine in mGluR-only system with MERC. **f**) Same as **e**) with no MERC. **g**) Calcium dynamics taken at 3 points in both the stimulated and neighboring spine in combined system with MERC. **h**) Same as **g**) with no MERC. **i**)IP_3_ dynamics taken at 3 points in both stimulated and neighboring spine in combined system with MERC. **j**) Same as **i**) with no MERC.

**Figure S3:**
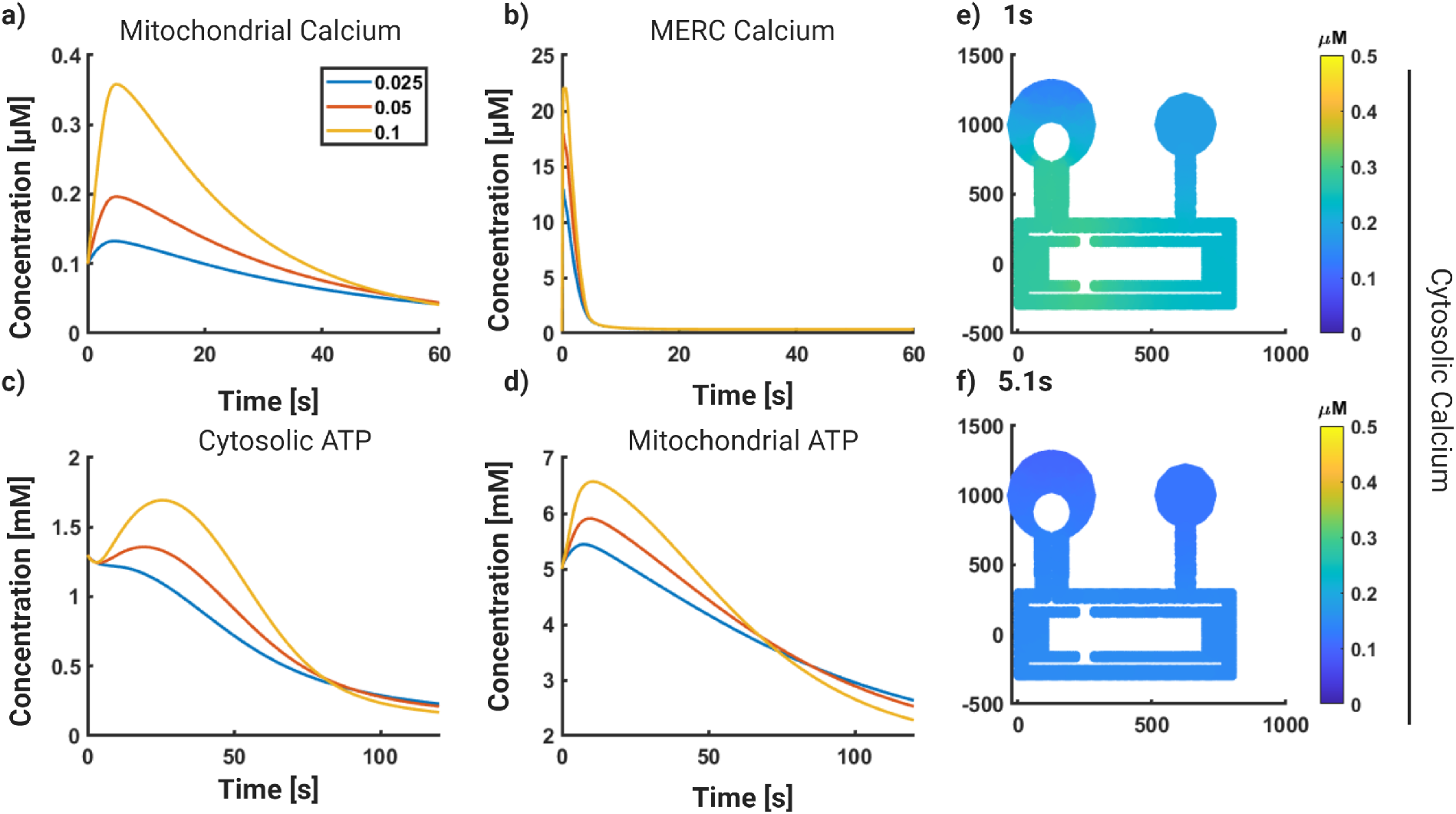
Simulations without glutamate response quickly decay toward steady-state. **a**)Mitochondrial calcium dynamics with no stimulus for MERC covering 2.5, 5, and 10 % mitochondria surface area. **b**) Same as **a**) for MERC Calcium. **c**) Same as **a**) for Cytosolic ATP. **d**) Same as **a**) for Mitochondrial ATP. **e**) 2D Spatial plot of cytosolic calcium with no stimulus at 1 second. **f**) same as **e**, but taken at 5.1s

**Figure S4:**
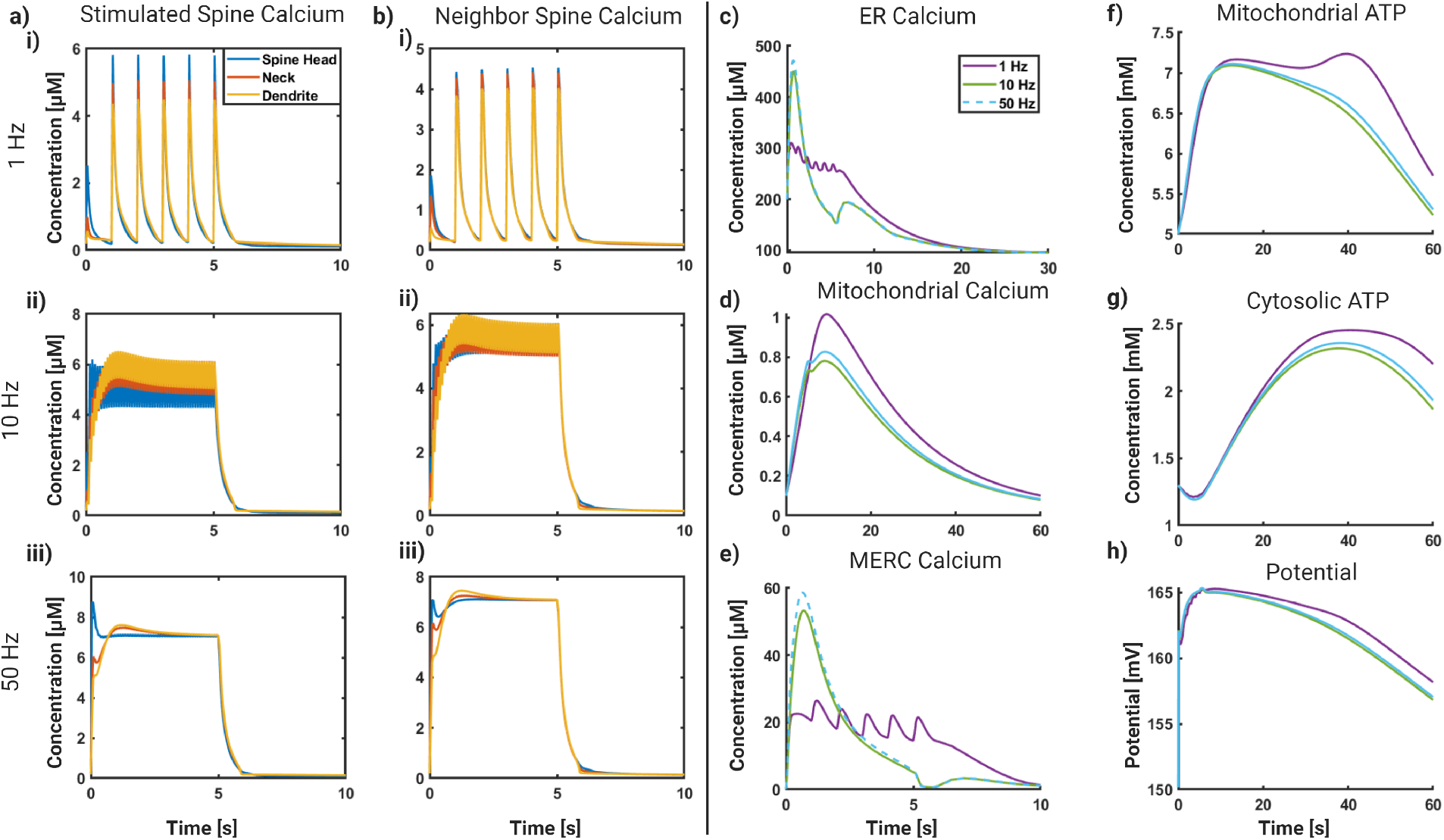
Changing stimulus frequency within range of expected signaling frequencies yields changes in system behavior. All results generated with mitochondria size of 0.6 μm and 10 percent MERC surface area. **a**) Stimulated spine calcium concentrations at 3 points in the spine cytosol for the following frequencies: **i**) 1 Hz **iib)**10 Hz **iiib)** 50 Hz **b**) Neighboring spine calcium concentrations at 3 points in the spine cytosols for ib) 1 Hz, **iib)** 10 Hz, **iiib)** 50 Hz. **c**) Volume average ER Calcium for the 3 frequencies. 10 Hz and 50 Hz frequency plots overlay. **d**) Same as **c**) for Mitochondrial Calcium. **e**) Same as **c**) for MERC Calcium. **f**) Same as **c**) for Mitochondrial ATP. **g**) Same as **c**) for Cytosolic ATP. **h**) Same as **c**) for Mitochondrial Potential

**Figure S5:**
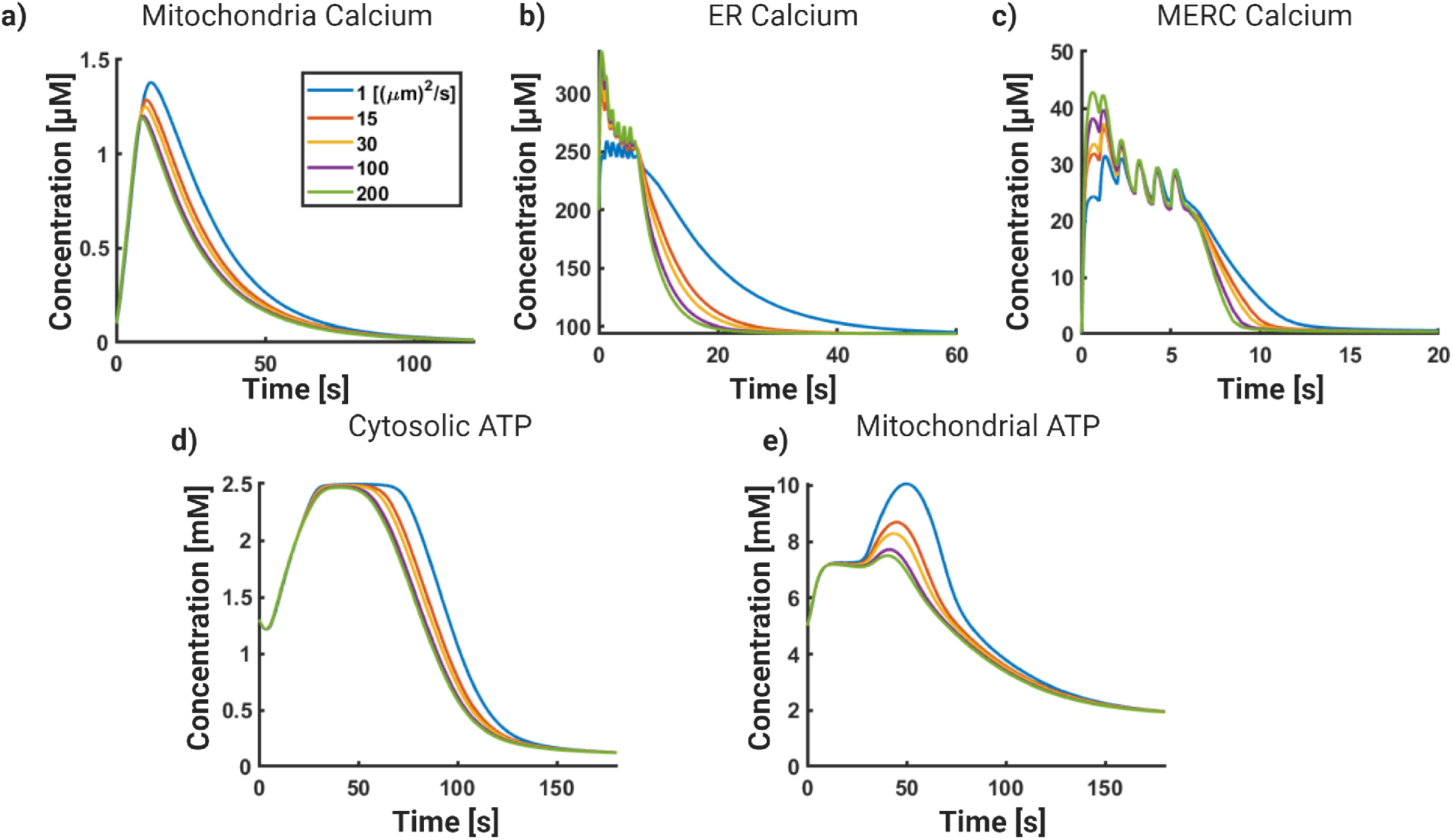
Calcium diffusion coefficient exhibits minor changes in system dynamics. **a**) Mitochondrial dynamics in response to 5 1 Hz pulses varying diffusion constants of free calcium. **b**) ER calcium dynamics with varying calcium diffusion constants. **c**) MERC calcium dynamics with varying calcium diffusion constants. **d**) Cytosolic ATP dynamics with varying calcium diffusion constants. **e**) Mitochondria ATP dynamics with varying calcium diffusion constants.

**Figure S6:**
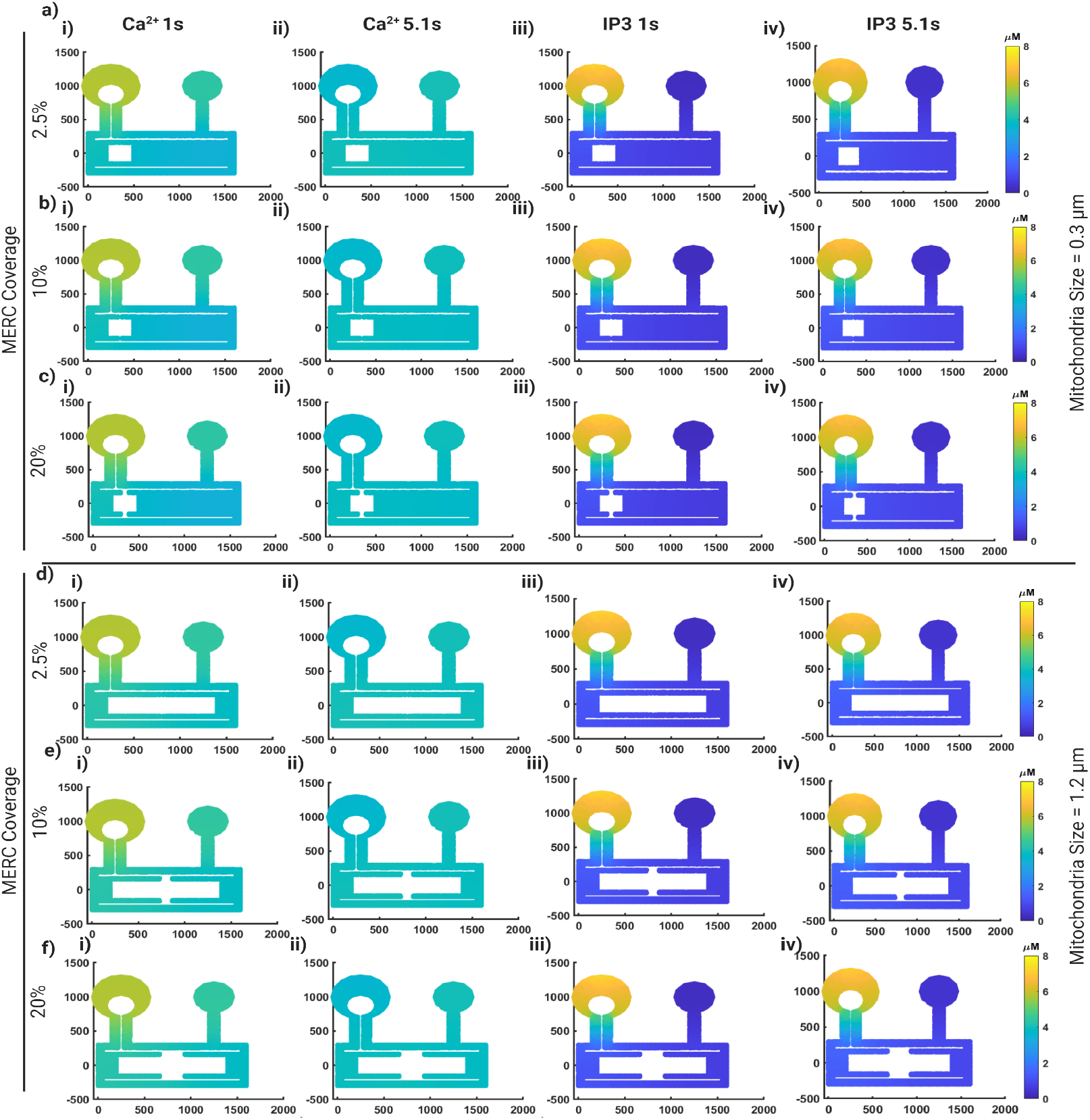
Cytosolic gradients do not change with changes in mitochondrial size or MERC surface area coverage. **a**) 2D Spatial plots for concentrations, in *μM*, for **i)** calcium at 1 second, **ii)** calcium at 5.1 seconds, **iii)** IP3 at 1 second, and **iv)** IP3 at 5.1 seconds for a mitochondrial size of 0.3 *μ*m and 2.5 % of mitochondria surface area covered by MERC. **b**) Same as **a**for a mitochondria size of 0.3 *μ*m and 10 % of mitochondria surface area covered by MERC. **c**) Same as **a**for a mitochondria size of 0.3 *μ*m and 20 % of mitochondria surface area covered by MERC. **d**) Same as **a**for a mitochondria size of 1.2 *μ*m and 2.5 % of mitochondria surface area covered by MERC. **e**) Same as **a**for a mitochondria size of 1.2 *μ*m and 10 % of mitochondria surface area covered by MERC. **f**) Same as **a**for a mitochondria size of 1.2 *μ*m and 20 % of mitochondria surface area covered by MERC.

**Figure S7:**
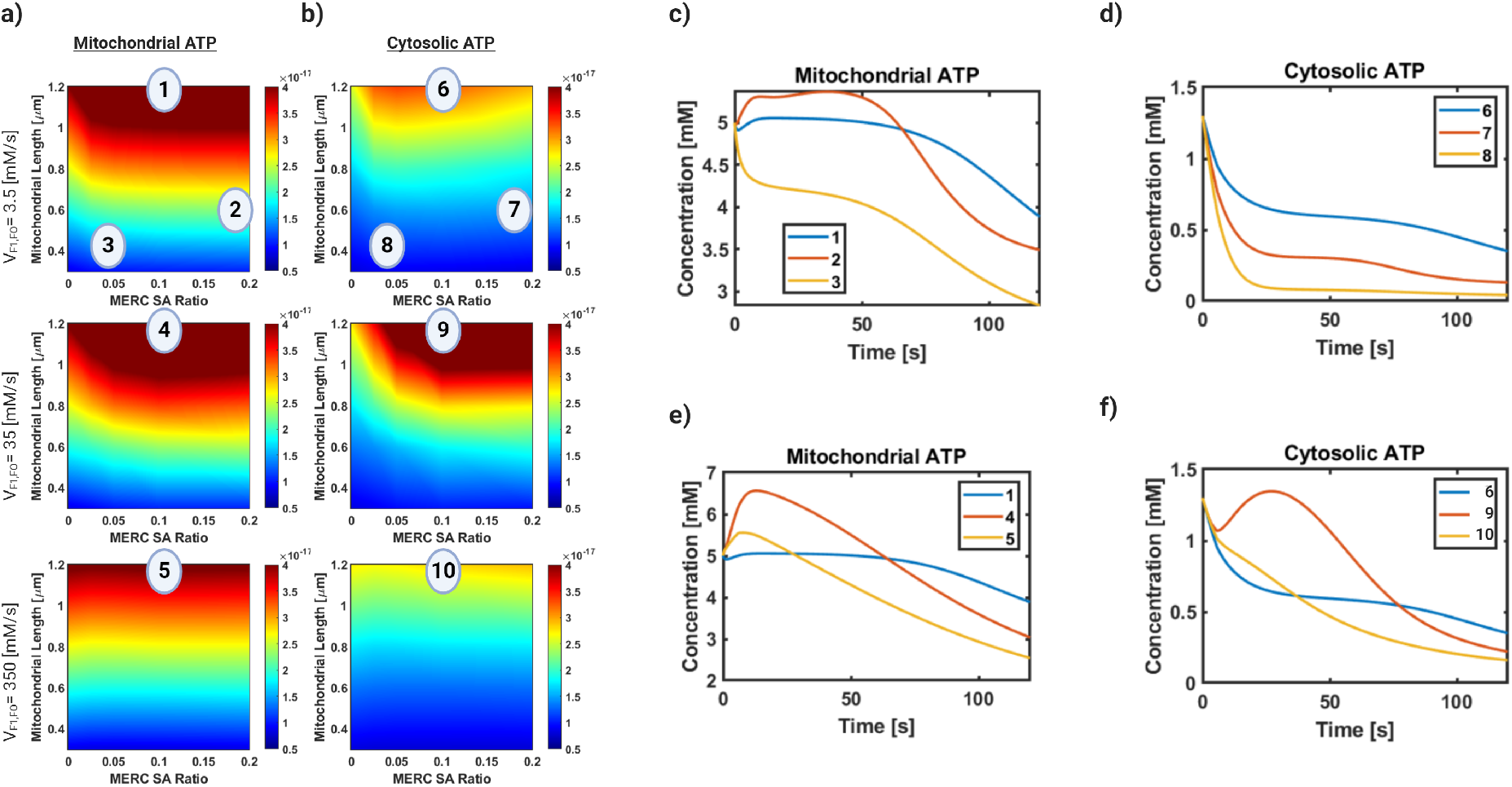
Dynamic plots for phase diagram show differential ATP generation. **a**) Phase diagrams for the change in V_*F*1*FO*_ parameter (shown in Figure 6) for mitochondrial ATP. Here we denote 5 points on the phase diagram and explore the corresponding dynamics in **c**) and **d**). **b**) Phase diagrams for the change in V_*F*1*FO*_ parameter (shown in Figure 6) for cytosolic ATP. Here we denote the same 5 points and explore the corresponding dynamics in **e**) and **f**). **c**) Comparison between 3 different phases displayed on the mitochondrial ATP phase diagram. **d**) Comparison between 3 different phases displayed on the cytosolic ATP phase diagram. The three dynamic plots behave similarly, but at different magnitudes. **e**) Comparison between the highest phases displayed on 3 different phase diagrams for mitochondrial ATP. Although all three diagrams have similar max AUC, higher V_*F*1*FO*_ values correspond to a higher initial production in this model. **f**) Comparison between the highest phases displayed on 3 different phase diagrams for mitochondrial ATP.

